# AKAP5 and Caveolin-1 organize opposing nanodomains that regulate smooth muscle contraction and blood pressure

**DOI:** 10.64898/2026.08.13.744495

**Authors:** Yen-Lin Chen, Maniselvan Kuppusamy, Fenix Araujo, Yashu Tang, Zdravka Daneva, Kyosuke Kazama, Lojy Hozyen, Emma D. Chung, Saainikedhana Venugopal, Sree S. Katragadda, Grace C. Garcia, Divine C. Nwafor, Stephen B. Abbott, Richard Minshall, Ryan T. Kellogg, Swapnil K. Sonkusare

## Abstract

TRPV4 ion channels in vascular smooth muscle cells (SMCs) are crucial regulators of blood pressure, and their functional effects are differentially shaped by their signaling partners. However, the mechanisms by which TRPV4 channels are compartmentalized into distinct signaling nanodomains with opposite impacts on blood pressure remain unclear. Here, we identify the scaffolding proteins that compartmentalize TRPV4 channels into discrete nanometer-scale signaling domains at the SMC plasma membrane and define how these nanodomains produce opposing effects on vasoconstriction and blood pressure. We show that AKAP5 anchors a nanodomain linking α1-adrenergic receptors, protein kinase C and TRPV4 channels, thereby driving sympathetic vasoconstriction and blood pressure elevation. In contrast, caveolin-1 promotes a mechanosensitive nanodomain comprising Piezo1, TRPV4, and BK channels that mediates vasodilation and a decrease in blood pressure. In hypertension, AKAP5-dependent constrictor nanodomains are hyperactive, whereas caveolin-1-based dilator nanodomains are hypoactive, shifting the balance toward pathological vasoconstriction. These findings reveal fundamental mechanisms that organize smooth muscle TRPV4 channels into spatially and functionally distinct nanodomains regulating blood pressure and show how disruption of this organization contributes to blood pressure elevation in hypertension.

## Introduction

Smooth muscle cells (SMCs) in small arteries play a crucial role in regulating vascular resistance, and therefore, blood pressure and tissue perfusion. SMC contraction is dynamically regulated by neurohumoral mediators and mechanical stimuli in the bloodstream. A variety of Ca^2+^ signaling mechanisms transduce these stimuli into SMC contraction, playing a critical role in maintaining vascular homeostasis (*1, 2*). Importantly, abnormal Ca^2+^ signaling in SMCs is a hallmark of hypertension (*1, 2*), highlighting the therapeutic potential of targeting these mechanisms to lower blood pressure.

Recent evidence underscores the importance of non-uniform protein distribution at the SMC plasma membrane in forming specialized nanodomains that confer signaling and functional specificity (*3–9*). Ca^2+^-permeable ion channels and their downstream targets are often colocalized within nanometer-scale domains at the plasma membrane, enabling precise and localized signaling (*3, 4, 7–9*). Transient receptor potential vanilloid 4 (TRPV4) channels are a crucial Ca^2+^ influx pathway in SMCs that regulates vascular resistance and blood pressure (*3*). Notably, TRPV4 channels increase SMC contraction and elevate blood pressure when positioned next to α1-adrenergic receptors (α1ARs), but promote relaxation and reduce blood pressure when adjacent to large-conductance Ca^2+^-activated K^+^ (BK) channels (*3, 10*). These findings support the concept that the functional impact of TRPV4 channels on SMC contraction is determined by the unique composition of their nanoscale environment (*9*). However, the molecular mechanisms that enable TRPV4 channels in SMCs to exert divergent functional effects through differential nanoscale organization remain poorly understood.

Scaffolding proteins play a central role in forming spatially localized signaling domains at the SMC plasma membrane. Caveolin-1 (Cav1) and A-kinase anchoring protein 5 (AKAP5) have been implicated in organizing ion channel networks within the vascular wall (*11–15*), regulating both vasoconstriction and vasodilation (*11, 14, 16, 17*). Both Cav1 and AKAP5 have also been shown to regulate TRPV4 channel activity in different cell types (*12, 15, 18*). Therefore, we hypothesized that spatially separate Cav1 and AKAP5 organize functionally distinct TRPV4 signaling nanodomains at the SMC membrane. Furthermore, we hypothesized that disruption of Cav1-and AKAP5-dependent TRPV4 signaling nanodomains contributes to enhanced vasoconstriction and elevated blood pressure in hypertension.

Our findings reveal that AKAP5 and Cav1 organize TRPV4 channel-containing Ca^2+^-signaling nanodomains at the SMC plasma membrane that exert functionally opposite effects. Sympathetic stimulation selectively activates AKAP5-dependent TRPV4 signaling via G-protein coupled receptors to raise blood pressure, whereas intraluminal pressure activates Cav1-based TRPV4 signaling through mechanosensitive Piezo1 channels to lower blood pressure. In hypertension, the activity of AKAP5-dependent constrictor nanodomains is increased, whereas the activity of Cav1-based dilator nanodomains is reduced, leading to increased vasoconstriction and elevated blood pressure. These findings reveal novel mechanisms by which anchoring proteins organize spatially separate TRPV4 channel-containing nanodomains at the SMC plasma membrane to produce distinct physiological and pathological effects and establish a framework for therapeutic intervention in hypertension.

## Results

### SMC AKAP5 (AKAP5_SMC_) promotes α1AR–protein kinase Cα–TRPV4 signaling to drive sympathetic vasoconstriction and blood pressure elevation

Ionic currents through TRPV4 channels were recorded in freshly isolated SMCs from third-order (∼100 μm diameter) mesenteric arteries (MAs) from male and female tamoxifen-fed AKAP5^fl/fl^ mice (controls) and SMC-AKAP5-knockout (AKAP5^SMCKO^) mice (tamoxifen-fed AKAP5^fl/fl^ *Myh11*-Cre^ERT2^RAD mice) using whole-cell patch-clamp electrophysiology (Figure 1A and Figure S1). Application of the α1AR agonist phenylephrine (1 μM) increased ionic currents in SMCs from control male and female mice that were inhibited by the TRPV4 inhibitor, GSK2193874 (100 nM; hereafter, GSK219). Phenylephrine-activated SMC TRPV4 (TRPV4_SMC_) currents were absent in AKAP5^SMCKO^ mice (Figure 1A and Figure S2A), suggesting that AKAP5_SMC_ is critical for α1AR–TRPV4_SMC_ signaling. However, direct activation of TRPV4 channels with the agonist GSK1016790A (100 nM; hereafter, GSK101) elicited similar TRPV4_SMC_ currents in control and AKAP5^SMCKO^ mice (Figure 1B), suggesting that AKAP5_SMC_ promotes α1AR–TRPV4_SMC_ signaling but does not alter the number of functional TRPV4_SMC_ channels at the cell membrane.

**Figure 1.**
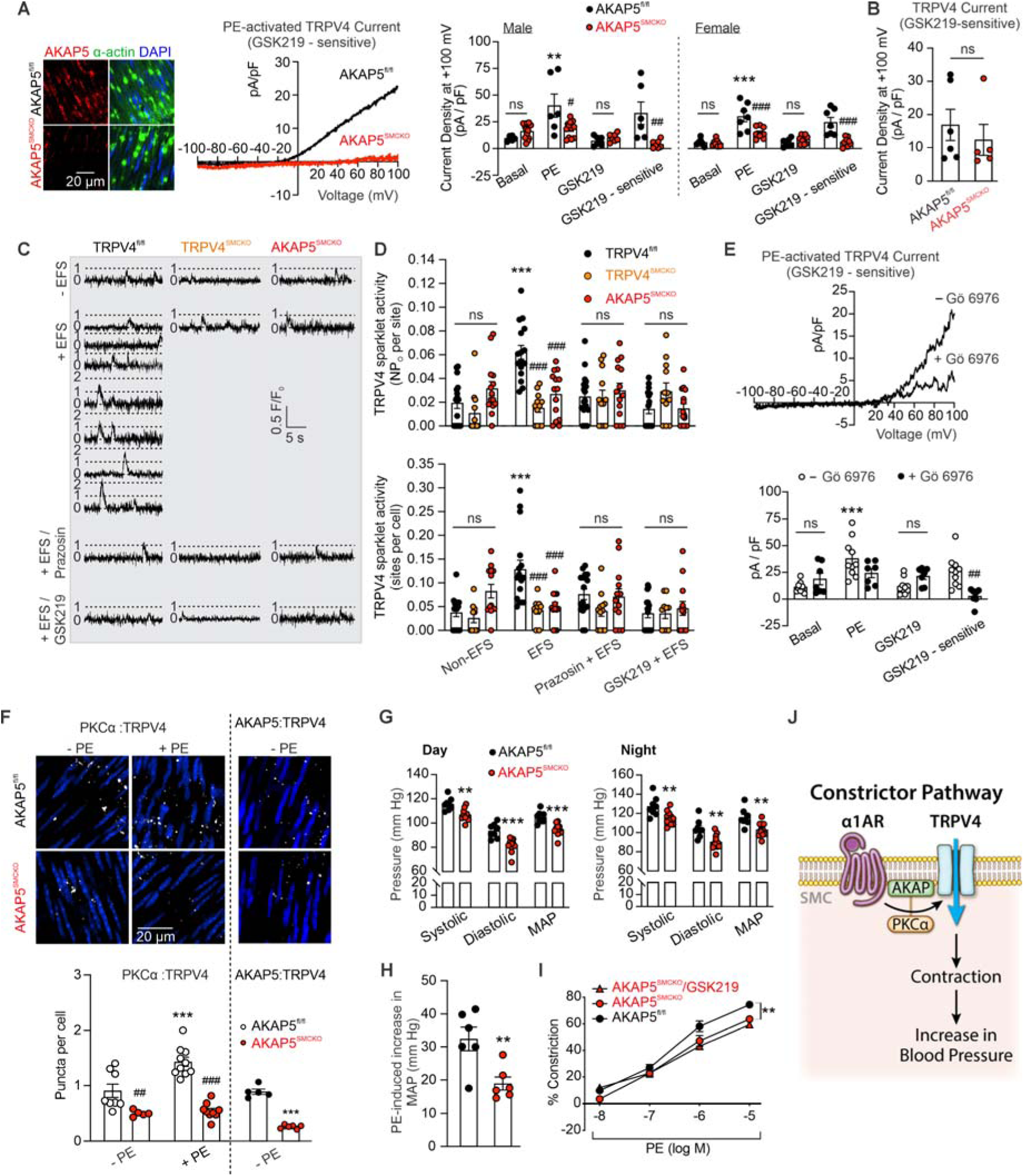
AKAP5_SMC_-dependent α1AR–PKCα–TRPV4_SMC_ signaling promotes sympathetic vasoconstriction and blood pressure elevation. (A) *Left*, representative images of AKAP5 (red), α-actin (green; SMC marker), and nuclear staining (DAPI; blue) in *en face* preparations of MAs from AKAP5^fl/fl^ and AKAP5^SMCKO^ mice. *Middle*, representative traces of ionic currents through TRPV4_SMC_ channels in freshly isolated SMCs from MAs of AKAP5^fl/fl^ and AKAP5^SMCKO^ mice, shown as phenylephrine (PE; 1 μM)-activated currents sensitive to the TRPV4 antagonist GSK219 (100 nM). Experiments were performed in the presence of ruthenium red (1 μM) to prevent Ca^2+^ entry at negative voltages and subsequent activation of BK channels. *Right*, averaged outward currents at +100 mV in isolated SMCs from MAs of AKAP5^fl/fl^ mice (male, n = 7; female, n = 7) and AKAP5^SMCKO^ mice (male, n = 8; female, n = 8) before and after treatment with PE (1 μM) or PE+GSK219 (**P < 0.01 vs. basal; ^#^P < 0.05; ^##^P < 0.01 vs. AKAP5^fl/fl^; ns, not significant; 2-way ANOVA). (B) Averaged GSK219-sensitive currents in isolated SMCs at +100 mV in the presence of the TRPV4 agonist GSK101 (100 nM) (n = 5; ns, not significant; unpaired t-test). (C) Representative fractional fluorescence (F/F_0_) traces reflecting sparklet activity in MAs from TRPV4^fl/fl^, TRPV4^SMCKO^, and AKAP5^SMCKO^ mice in response to EFS. Prior to EFS, applied as 5-second trains of electrical pulses (50–150 V, 0.25-ms, 15 Hz), Fluo-4AM–loaded, pressurized MAs were pretreated with cyclopiazonic acid (CPA; 20 μM), to prevent intracellular Ca^2+^-release signals; capsaicin (1 μM), to desensitize sensory nerves; atropine (10 μM), to block muscarinic receptors, and nifedipine (1 μM); to block L-type Ca^2+^ channels. Experiments were performed in the absence or presence of prazosin (1 μM; α1AR antagonist) or GSK219 (100 nM; TRPV4 inhibitor). Dotted lines represent quantal levels (single-channel amplitudes) determined from all-points histograms. (D) *Upper*, averaged TRPV4 sparklet activity per site (NP_O_, where N is the number of channels and P_O_ is the open state probability) and (*lower*) TRPV4 sparklet sites per cell in pressurized MAs from TRPV4^fl/fl^ (n = 16), TRPV4^SMCKO^ (n = 12), and AKAP5^SMCKO^ (n = 14) mice under basal conditions (CPA/capsaicin/atropine/nifedipine) and in response to EFS in the absence or presence of prazosin (1 μM) or GSK219 (100 nM) (***P < 0.001 vs. Basal; ^###^P < 0.001 vs.TRPV4^fl/fl^; ns, not significant; 2-way ANOVA). (E) *Upper*, representative traces of ionic currents through TRPV4 channels in freshly isolated SMCs from MAs of AKAP5^fl/fl^ mice in the absence or presence of Gö 6976 (PKCα/β inhibitor, 1 μM), shown as as PE (1 μM)-activated, GSK219 (100 nM)-sensitive currents. Experiments were performed in the presence of RuR (1 μM) to block Ca^2+^ entry at negative voltages. *Lower*, outward currents at +100 mV in SMCs from MAs of AKAP5^fl/fl^ mice before and after administration of PE (1 μM) or PE+GSK219 (100 nM) in the absence (n = 7) or presence (n = 9) of Gö 6976 (1 μM) (***P < 0.001 vs. basal; ^##^P < 0.01 vs. Gö 6976; ns, not significant; 2-way ANOVA). (F) *Upper*, representative *in situ* PLA images showing SMC nuclei (blue) and PKCα:TRPV4_SMC_ colocalization (white puncta) before and after PE treatment, and AKAP5_SMC_:TRPV4_SMC_ colocalization (white puncta) in the absence of PE in *en face* preparations of MAs from AKAP5^fl/fl^ and AKAP5^SMCKO^ mice. *Lower*, quantification of PKCα:TRPV4_SMC_ colocalization in MAs from AKAP5^fl/fl^ and AKAP5^SMCKO^ mice before (AKAP5^fl/fl^, n = 8; AKAP5^SMCKO^, n = 5) and after (AKAP5^fl/fl^, n = 10; AKAP5^SMCKO^, n = 9) PE treatment (***P < 0.001 vs. without PE; ^##^P < 0.01, ^###^P < 0.001 vs. AKAP5^fl/fl^; 2-way ANOVA). Quantification of AKAP5_SMC_:TRPV4_SMC_ colocalization in MAs from AKAP5^fl/f^ (n = 5) and AKAP5^SMCKO^ (n = 6) mice (***P < 0.001 vs. AKAP5^fl/fl^; unpaired t test). (G) Quantification of systolic and diastolic blood pressure and mean arterial pressure (MAP) during daytime (left) and nighttime (right) in AKAP5^fl/fl^ (n = 8) and AKAP5^SMCKO^ (n = 11) mice (**P < 0.01, ***P < 0.001 vs. AKAP5^fl/fl^; 2-way ANOVA). (H) Increases in MAP after a single bolus injection of PE (10 mg/kg, ip) in AKAP5^fl/fl^ (n = 6) and AKAP5^SMCKO^ (n = 6) mice (**P < 0.01 vs. AKAP5^fl/fl^; unpaired t test). (I) Pressure myography data showing PE-induced constriction of MAs in the absence or presence of the TRPV4 inhibitor GSK219 (100 nM) (AKAP5^fl/fl^, n = 7; AKAP5^SMCKO^, n = 5; AKAP5^SMCKO^+GSK219, n = 6; **P < 0.01 vs. AKAP5^fl/fl^; 2-way ANOVA). (J) Schematic depicting AKAP5_SMC_-dependent α1AR–PKCα–TRPV4_SMC_ signaling for sympathetic vasoconstriction and blood pressure elevation.

To determine whether sympathetic nerve stimulation activates TRPV4_SMC_ channels through α1AR, we used electrical field stimulation (EFS), applying 5-second trains of electrical pulses (50–150 V, 0.25-ms pulse, 15 Hz) in the presence of capsaicin (1 μM; to densitize sensory nerves), atropine (10 μM; muscarinic receptor antagonist), and nifedipine (1 μM; L-type Ca^2+^ channel antagonist) in cannulated and pressurized MAs (intraluminal pressure, 80 mm Hg). Nerve stimulation increased the activity of elementary Ca^2+^-influx signals through TRPV4_SMC_ channels (TRPV4 sparklets) (*19*) in control mice; this effect was abolished in the presence of the α1AR antagonist prazosin (1 μM), suggesting activation of α1AR–TRPV4_SMC_ signaling by nerve stimulation. Nerve-stimulation–induced increases in TRPV4_SMC_ sparklet activity were absent in MAs from TRPV4^SMCKO^ and AKAP5^SMCKO^ mice (Figure 1C, D), providing evidence for a critical role of AKAP5_SMC_ in nerve-stimulation–induced α1AR–TRPV4_SMC_ signaling.

AKAP5-anchoring of protein kinase Cα (PKCα) has been shown to promote TRPV4 phosphorylation and activation in different cell types (*8, 18, 20, 21*). Therefore, we tested whether AKAP5_SMC_ mediates α1AR– PKCα–TRPV4_SMC_ signaling. Whole-cell patch-clamp studies in freshly isolated SMCs from control mice showed that α1AR-mediated activation of TRPV4_SMC_ channels was reduced by the PKCα inhibitor, Gö 6976 (1 μM) (*22*) (Figure 1E), supporting α1AR–PKCα–TRPV4_SMC_ signaling. Additionally, an *in situ* proximity ligation assay (PLA), which detects two proteins within ∼40 nm from one another, revealed that TRPV4_SMC_ channels colocalize with both PKCα and AKAP5_SMC_ under basal conditions. PKCα:TRPV4_SMC_ colocalization was significantly reduced in AKAP5^SMCKO^ mice, suggesting that AKAP5_SMC_ promotes the proximity of PKCα with TRPV4_SMC_ channels. Notably, α1AR stimulation for 5 minutes increased PKCα:TRPV4_SMC_ colocalization in MAs from control mice but not in those from AKAP5^SMCKO^ mice (Figure 1F). Together, these results suggest that AKAP5_SMC_ is critical for PKCα:TRPV4_SMC_ colocalization and α1AR–PKCα–TRPV4_SMC_ signaling.

Daytime and nighttime systolic, diastolic, and mean arterial pressures, recorded using radiotelemetry, were lower in AKAP5^SMCKO^ mice compared with AKAP5^fl/fl^ control mice (Figure 1G and Figure S3A); however, heart rate was not different between genotypes (Figure S3B, C). An acute injection of the α1AR agonist, phenylephrine (10 mg/kg, intraperitoneally [ip]), increased blood pressure in control mice; this effect was significantly reduced in AKAP5^SMCKO^ mice (Figure 1H and Figure S3D), indicating a crucial role for AKAP5_SMC_ in α1AR-induced elevation of blood pressure.

In pressure myography experiments, the TRPV4 inhibitor, GSK219 (100 nM), inhibited constriction of MAs in response to phenylephrine (10 nM to 10 μM) (*3*). Phenylephrine-induced constriction of MAs was also attenuated in MAs from AKAP5^SMCKO^ mice (Figure 1I), revealing the importance of both AKAP5_SMC_ and TRPV4_SMC_ channels in α1AR-mediated vasoconstriction. Moreover, the TRPV4 inhibitor GSK219 did not cause a further decrease in phenylephrine-induced constriction in AKAP5^SMCKO^ mice (Figure 1I), suggesting that AKAP5_SMC_ primarily mediates the TRPV4 component of phenylephrine-induced constriction. Overall, these data provide evidence that AKAP5 promotes constrictor α1AR–PKCα–TRPV4_SMC_ signaling, which elevates blood pressure (Figure 1J).

### Cav1 facilitates dilator TRPV4_SMC_–BK signaling but not constrictor **α**1AR–TRPV4 signaling, whereas AKAP150_SMC_ promotes **α**1AR–TRPV4 signaling but not TRPV4_SMC_–BK signaling

In direct contrast to the case for AKAP5^SMCKO^ mice (Figure 1G), resting systolic, diastolic, and mean arterial pressure were elevated in Cav1^SMCKO^ mice compared with Cav1^fl/fl^ control mice (Figures 2A, B), suggesting that Cav1_SMC_ lowers resting blood pressure. The increase in resting blood pressure in Cav1^SMCKO^ mice was observed in both daytime and nighttime recordings (Figure 2A), and was not associated with a change in resting heart rate (Figure S4). Consistent with these data, endothelium-denuded MAs from Cav1^SMCKO^ mice showed higher pressure-induced (myogenic) constriction than MAs from Cav1^fl/fl^ mice (Figure 2C), suggesting that a dilatory effect of Cav1_SMC_ limits myogenic constriction of small arteries.

**Figure 2.**
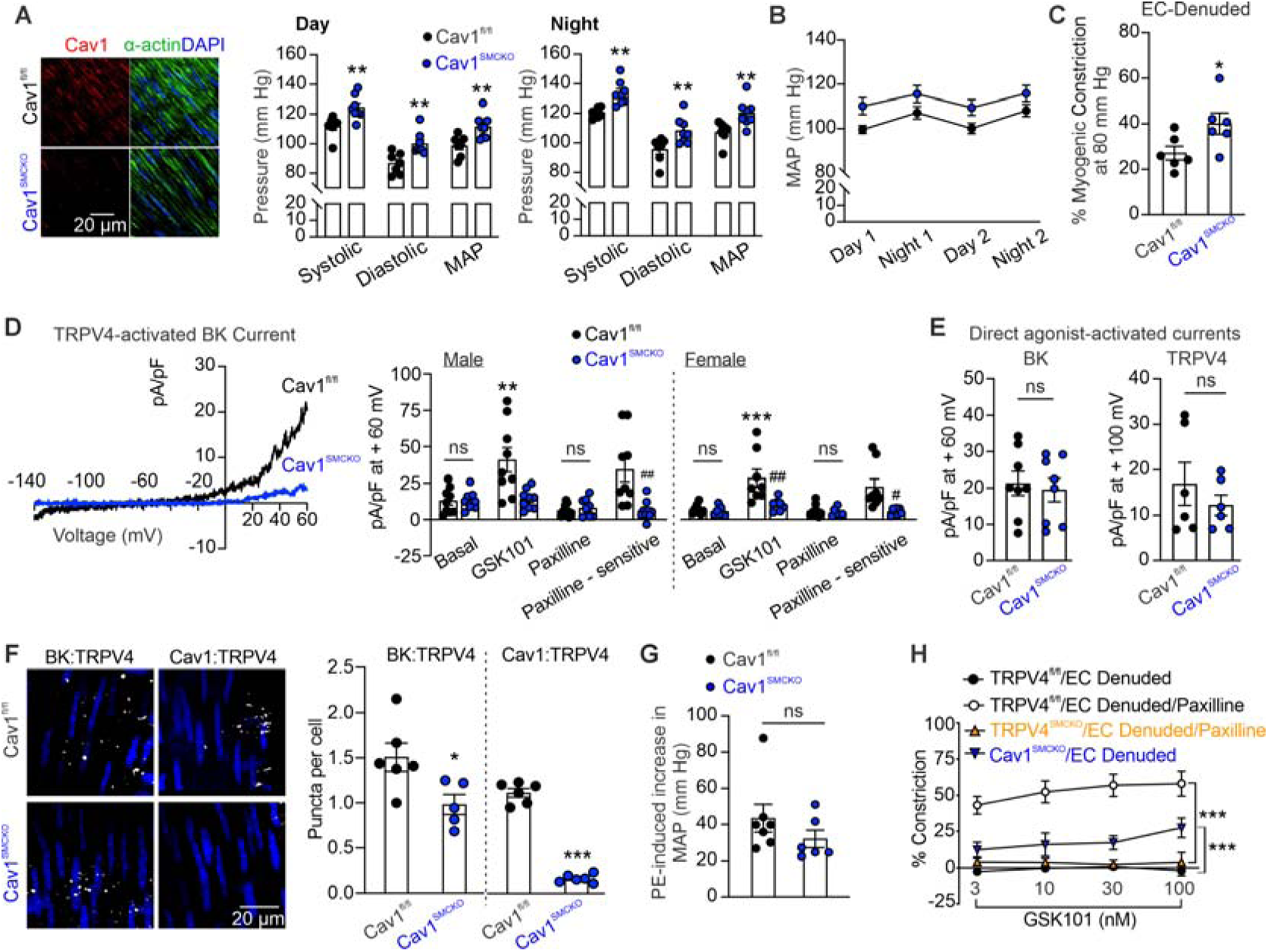
Cav1_SMC_ promotes blood-pressure–lowering dilator TRPV4_SMC_–BK signaling. (A) *Left*, representative images of Cav1 (red), α-actin (green; SMC marker), and nuclear staining (DAPI; blue) in *en face* preparations of MAs from Cav1^fl/fl^ and Cav1^SMCKO^ mice. Quantification of systolic and diastolic blood pressure and mean arterial pressure (MAP) during daytime (*middle*) and nighttime (*right*) in Cav1^fl/fl^ (n = 7) and Cav1^SMCKO^ (n = 8) mice (**P < 0.01 vs. Cav1^fl/fl^; 2-way ANOVA). (B) Resting MAP in Cav1^fl/fl^ (n = 7) and Cav1^SMCKO^ (n = 8) mice recorded over 2 days and 2 nights. (C) Averaged data showing the development of myogenic tone at 80 mm Hg intraluminal pressure in EC-denuded MAs from Cav1^fl/fl^ (n = 6) and Cav1^SMCKO^ (n = 6) mice (*P < 0.05 vs. Cav1^fl/fl^; unpaired t-test). (D) *Left*, representative traces of ionic currents through BK channels in freshly isolated SMCs from MAs of Cav1^fl/fl^ and Cav1^SMCKO^ mice. *Right*, outward currents at +60 mV in SMCs from MAs of Cav1^fl/fl^ (male, n = 9; female, n = 8) and Cav1^SMCKO^ (male, n = 9; female, n = 7) mice before and after GSK101 (TRPV4 agonist, 30 nM) or GSK101+paxilline (BK inhibitor, 1 μM) (**P < 0.01, ***P < 0.001 vs. basal; ^#^P < 0.05, ^##^P < 0.01 vs. Cav1^fl/fl^; ns, not significant; 2-way ANOVA). (E) *Left*, GoSlo SR 5-69 (BK channel activator; 1 μM)–activated outward currents (GoSlo SR 5-69 minus baseline current) at +60 mV in SMCs isolated from MAs of Cav1^fl/fl^ (n = 8) and Cav1^SMCKO^(n = 8) mice (ns, not significant; unpaired t-test). *Right*, GSK101 (TRPV4 agonist, 100 nM)-activated, GSK219 (100 nM)–sensitive outward currents at +60 mV in SMCs from MAs of Cav1^fl/fl^ (n = 6) and Cav1^SMCKO^ mice (n = 6) (ns, not significant; unpaired t-test). (F) *Left*, representative PLA images showing SMC nuclei (blue) and BK:TRPV4_SMC_, and Cav1_SMC_:TRPV4_SMC_ colocalization (white puncta) in *en face* preparations of MAs from Cav1^fl/fl^ and Cav1^SMCKO^ mice. *Right*, quantification of BK:TRPV4_SMC_ and Cav1_SMC_:TRPV4_SMC_ colocalization in MAs from Cav1^fl/fl^ (n = 6) and Cav1^SMCKO^ (n = 6) mice (*P < 0.05, ***P < 0.001 vs. Cav1^fl/fl^; unpaired t-test). (G) Increases in MAP after a single bolus injection of PE (10 mg/kg, ip) in Cav1^fl/fl^ (n = 7) and Cav1^SMCKO^ (n = 6) mice (unpaired t-test). (H) Pressure myography data showing GSK101-induced constriction of EC-denuded MAs in the absence or presence of the BK channel antagonist paxilline (1 μM; TRPV4^fl/fl^, n = 11; TRPV4^fl/fl^/paxilline, n = 7; TRPV4^SMCKO^/paxilline, n = 7; Cav1^SMCKO^, n = 8; ***P < 0.001 vs. TRPV4^fl/fl^; 2-way ANOVA).

We then tested the possibility that Cav1_SMC_, but not AKAP5_SMC_, promotes dilator TRPV4_SMC_–BK channel signaling. Patch-clamp studies on freshly isolated SMCs from male and female mice showed that the TRPV4 agonist GSK101 activates outward BK currents that are inhibited by the BK channel inhibitor paxilline (1 μM). TRPV4_SMC_-activated BK currents were reduced in SMCs from Cav1^SMCKO^ mice (Figure 2D and Figure S2C), suggesting a critical role for Cav1_SMC_ in TRPV4_SMC_–BK channel signaling. BK currents directly activated by the BK channel agonist GoSlo SR 5-69 (1 μM) (Figure 2E), and TRPV4 currents elicited by the TRPV4 agonist GSK101 (100 nM) (Figure 2E), were not different between SMCs from Cav1^fl/fl^ and Cav1^SMCKO^ mice, supporting an impairment in TRPV4_SMC_–BK channel signaling in Cav1^SMCKO^ mice rather than a decrease in the number of functional TRPV4 or BK channels at the cell membrane. A proximity analysis (PLA) suggested that TRPV4_SMC_ channels colocalize with Cav1 and BK channels in MAs. We further found that TRPV4:BK colocalization was reduced in SMCs from Cav1^SMCKO^ mice (Figure 2F), supporting a critical role for Cav1_SMC_ in TRPV4_SMC_:BK colocalization and TRPV4_SMC_–BK channel signaling.

TRPV4-induced increases in BK current were not altered in SMCs from AKAP5^SMCKO^ mice (Figure S2B and Figure S5A, B), suggesting that AKAP5_SMC_ does not play a role in dilator TRPV4_SMC_–BK signaling. Moreover, phenylephrine-induced increases in TRPV4_SMC_ channel currents were not different between SMCs from control and Cav1^SMCKO^ mice (Figure S2D & S6), and phenylephrine (10 mg/kg, ip)-induced increases in blood pressure were not altered in Cav1^SMCKO^ mice (Figure 2G), indicating that Cav1_SMC_ does not regulate constrictor α1AR– TRPV4_SMC_ signaling.

To functionally separate constrictor and dilator TRPV4_SMC_ subpopulations, we treated endothelial cell (EC)-denuded MAs with the TRPV4 agonist, GSK101 (3–100 nM). GSK101 did not alter the diameter of MAs from control mice, likely due to simultaneous activation of both constrictor and dilator TRPV4_SMC_ channels. However, inhibiting the dilator nanodomains with the BK channel inhibitor paxilline (1 μM) unmasked a large vasoconstrictor response to GSK101 (Figure 2H). Moreover, TRPV4_SMC_-mediated vasoconstriction was increased in Cav1^SMCKO^ mice and was absent in MAs from TRPV4^SMCKO^ mice (Figure 2H). These data provide functional evidence for separate constrictor and dilator TRPV4_SMC_ channel signaling in MAs.

### Deletion of AKAP5_SMC_ or Cav1_SMC_ disrupts constrictor or dilator TRPV4_SMC_ nanodomains, respectively

To determine the nanoscale SMC plasma membrane localization of TRPV4_SMC_ with PKCα and BK channels, used as markers for constrictor and dilator nanodomains, respectively, we used simultaneous multicolor total internal reflection fluorescence single-molecule localization microscopy (TIRF-SMLM; 15 nm resolution), with application of the spectral demixing principle (Figure S7) (*23*), which eliminates chromatic aberration issues and drift associated with sequential color acquisitions. Localization maps for TRPV4_SMC_, PKCα, and BK channels were generated in freshly isolated SMCs from MAs (Figure 3A). A colocalization analysis showed that ∼8% of total TRPV4_SMC_ channels colocalized (occurred within 50 nm or less) with PKCα, whereas ∼9% of TRPV4_SMC_ channels colocalized with BK channels. TRPV4_SMC_:PKCα colocalization was significantly reduced in SMCs from AKAP5^SMCKO^ mice compared to control mice, but was unchanged in SMCs from Cav1^SMCKO^ mice. Conversely, TRPV4_SMC_:BK colocalization was reduced in SMCs from Cav1^SMCKO^ mice compared to control mice, but was not altered in SMCs from AKAP5^SMCKO^ mice (Figure 3B). Deletion of AKAP5_SMC_ or Cav1_SMC_ did not alter the molecular density (Figure 3C) or cluster density (Figure S8) of TRPV4_SMC_, PKCα or BK channels, indicating that the scaffolding proteins do not alter the total number of proteins at the cell membrane. Moreover, triple-colocalization analysis of TRPV4_SMC_, BK, and PKCα showed that only ∼ 1% of TRPV4_SMC_ channels colocalized with both PKCα and BK channels, and this triple-colocalization was unaltered in SMCs from AKAP5^SMCKO^ or Cav1^SMCKO^ mice (Figure 3C). These data support the concept that AKAP5_SMC_ and Cav1_SMC_ facilitate the formation of spatially separate constrictor and dilator TRPV4_SMC_-signaling nanodomains, respectively (Figure 3D).

**Figure 3.**
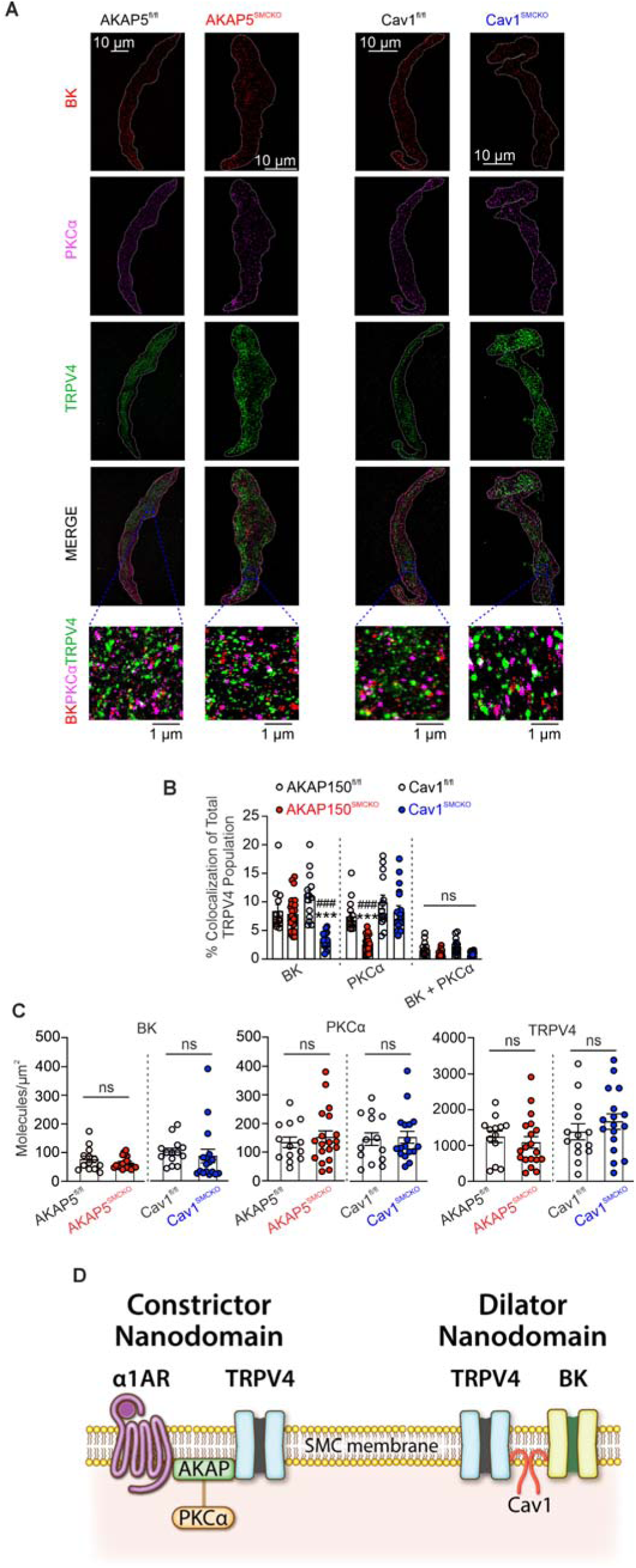
AKAP5_SMC_ and Cav1_SMC_ promote constrictor and dilator TRPV4_SMC_ signaling, respectively. (A) Representative TIRF-SMLM images showing localization maps for BK, PKCα, and TRPV4_SMC_ channels at the cell membrane of SMCs from MAs of AKAP5^fl/fl^, AKAP5^SMCKO^, Cav1^fl/fl^, and Cav1^SMCKO^ mice. Images were acquired with simultaneous multicolor TIRF-SMLM (15-nm resolution) using a spectral demixing principle. (B) Summary of colocalization analyses showing the percentage of total TRPV4_SMC_ channels within 50 nm of BK channels or PKCα or both in SMCs from AKAP5^fl/fl^ (n = 8), AKAP5^SMCKO^ (n = 10), Cav1^fl/fl^ (n = 8), and Cav1^SMCKO^ (n = 10) mice (1–2 cells/mouse; *P < 0.05, ***P < 0.001 vs. AKAP5^fl/fl^; ^##^P < 0.01, ^###^P < 0.001 vs. Cav1^fl/fl^; 2-way ANOVA). (C) Summary of dSTORM TIRF-SMLM superresolution localization maps showing the molecular density of BK, PKCα, and TRPV4_SMC_ channels at the plasma membrane of SMCs from AKAP5^fl/fl^ (n = 8), AKAP5^SMCKO^ (n = 10), Cav1^fl/fl^ (n = 8), and Cav1^SMCKO^ (n = 10) mice (1–2 cells/mouse; ns, not significant; unpaired t-test). (D) Schematic depicting the spatially separated AKAP5-dependent PKCα– TRPV4_SMC_ and Cav1-based TRPV4_SMC_–BK signaling nanodomains.

### Piezo1 channels impart mechanosensitivity to dilator TRPV4_SMC_–BK nanodomains

Dilator TRPV4_SMC_–BK signaling reduced vasoconstriction in response to pressure—a mechanical stimulus (Figure 2C, D). However, whether TRPV4 channels are genuinely mechanosensitive has been disputed (*24*), raising the possibility that an upstream mechanosensor may mediate pressure-induced activation of dilator TRPV4_SMC_ channels. Accordingly, we hypothesized that mechanosensitive Piezo1 channels, which are present in SMCs, mediate pressure-induced activation of dilator TRPV4_SMC_–BK channel signaling.

Patch-clamp recordings in freshly isolated SMCs from control mice showed that mechanical activation of cell-attached patches, achieved by calibrated negative pressure stimuli (−20 to -60 mm Hg), increased inward currents (Figure 4A), with a quantal level (unitary amplitude) of -2.4 pA (Figure S9A, B). Notably, mechanically activated inward currents were absent in SMCs from Piezo1^SMCKO^ mice (Figure 4A). The Piezo1 agonist, Yoda1 (10 μM), also activated inward currents that were reduced in SMCs from Piezo1^SMCKO^ mice (Figure 4B), providing evidence that Piezo1 currents are present in SMCs from MAs. Piezo1 currents, inhibited by the non-selective Piezo1 inhibitor GsMTx4 (5 μM), were also observed in freshly isolated SMCs from human skeletal muscle artery SMCs (Figure S10).

**Figure 4.**
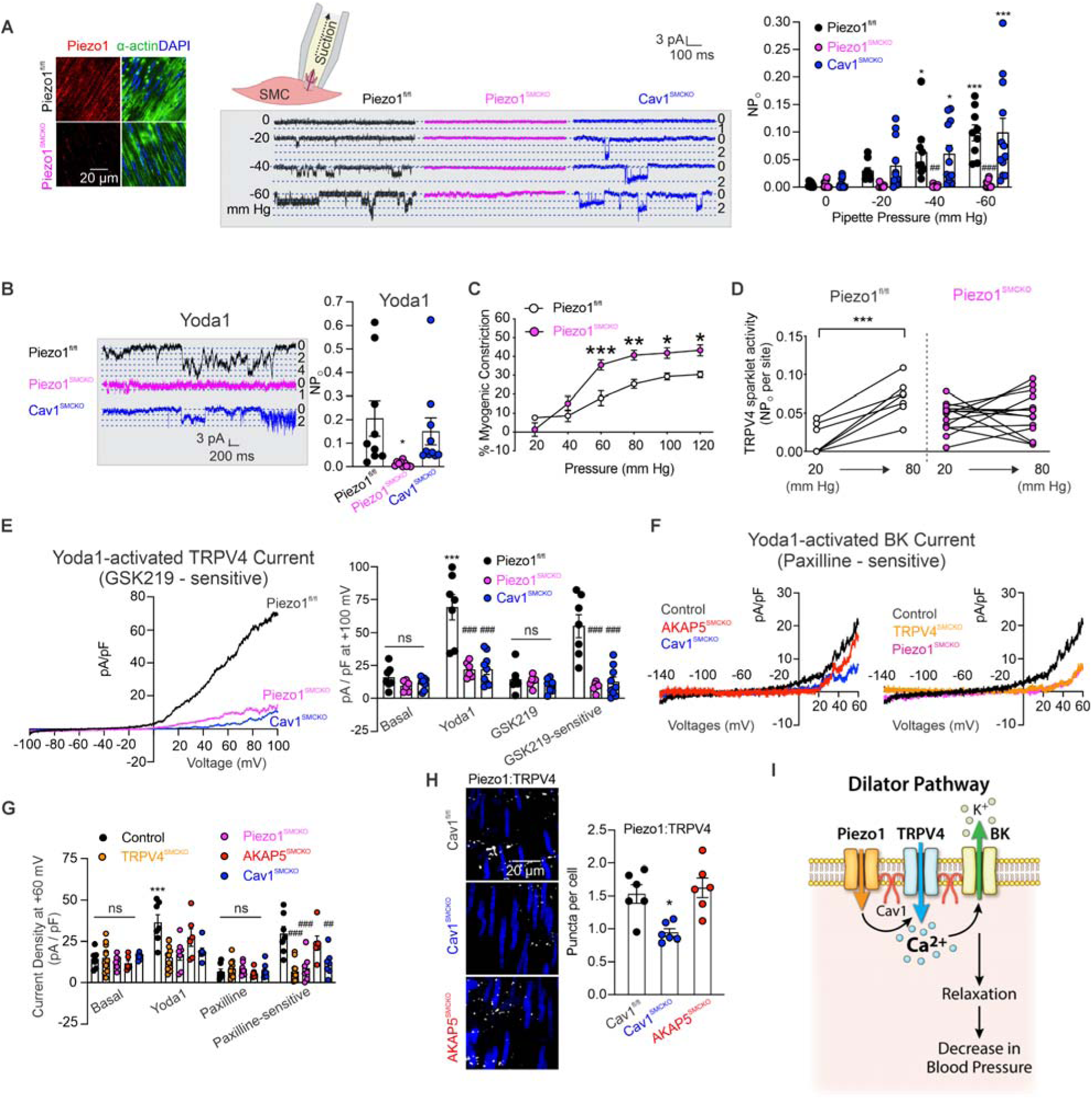
Piezo1 channels impart mechanosensitivity to Cav1-dependent TRPV4_SMC_–BK nanodomains. (A) *Left*, representative images of Piezo1 (red), α-actin (green; SMC marker), and nuclear staining (DAPI; blue) in *en face* preparations of MAs from Piezo1^fl/fl^ and Piezo1^SMCKO^ mice. *Inset*, schematic diagram showing suction applied to a SMC through a patch pipette for single-channel recording of Piezo1 currents. *Middle,* representative traces of stretch-activated inward currents in freshly isolated SMCs from MAs of Piezo1^fl/fl^, Piezo1^SMCKO^, and Cav1^SMCKO^ mice. Gradually increasing suction (0, -20, -40, and -60 mm Hg) elicited inward currents. Dotted lines represent quantal levels (single-channel amplitudes) determined from all-points histograms (Figure S9). *Right*, Piezo1 channel activity (NP_O_, where N is the number of channels and P_O_ is the open state probability) with gradually increased suction in SMCs from Piezo1^fl/fl^ (n = 9), Piezo1^SMCKO^ (n = 16), and Cav1^SMCKO^ (n = 12) mice (*P < 0.05, ***P < 0.001 vs. 0 mm Hg pipette pressure; ^#^P < 0.05, ^##^P < 0.01, ^###^P < 0.001 vs. Piezo1^fl/fl^; 2-way ANOVA). (B) *Left*, representative traces of currents induced by the Piezo1 channel activator Yoda1 (10 μM) in isolated SMCs from Piezo1^fl/fl^, Piezo1^SMCKO^, and Cav1^SMCKO^ mice. *Right*, Piezo1 channel activity (NP_O_) with Yoda1 (10 μM) in SMCs from Piezo1^fl/fl^ (n = 9), Piezo1^SMCKO^ (n = 10), and Cav1^SMCKO^ (n = 10) mice (*P < 0.05 vs. Piezo1^fl/fl^; 1-way ANOVA). (C) Averaged data showing the myogenic tone of MAs in response to different intraluminal pressures in Piezo1^fl/fl^ (n = 7) and Piezo1^SMCKO^ (n = 8) mice (*P < 0.05, **P < 0.01, ***P < 0.001 vs. Piezo1^fl/fl^; 2-way ANOVA). (D) TRPV4_SMC_ sparklet activity (NP_O_ per site) in MAs from Piezo1^fl/fl^ (n = 7) and Piezo1^SMCKO^ (n = 14) mice. Pressurized MAs were pretreated with cyclopiazonic acid (CPA; 20 μM) to eliminate interference from intracellular Ca^2+^ release, nifedipine (1 μM) to block L-type Ca^2+^ channels and GSK101 (30 nM) to stimulate TRPV4 agonist, and intraluminal pressure was increased from 20 mm Hg to 80 mm Hg (***P < 0.001 vs. 20 mm Hg; paired t-test). (E) *Left*, representative traces of Yoda1-induced ionic currents through TRPV4_SMC_ channels in SMCs from MAs of Piezo1^fl/fl^, Piezo1^SMCKO^ and Cav1^SMCKO^ mice, presented as GSK219-sensitive currents. *Right*, outward currents at +100 mV in SMCs from MAs of Piezo1^fl/fl^ (n = 7), Piezo1^SMCKO^ (n = 8), and Cav1^SMCKO^ (n = 8) mice before and after addition of Yoda1 (10 μM) or Yoda1+GSK219 (100 nM) (***P < 0.001 vs. basal; ^###^P < 0.001 vs. Piezo1^fl/fl^; ns, not significant; 2-way ANOVA). (F) Representative traces of Yoda1-induced BK channel currents in SMCs from MAs of Control, AKAP5^SMCKO^, Cav1^SMCKO^, Piezo1^SMCKO^ and TRPV4^SMCKO^ mice, presented as paxilline (BK inhibitor)-sensitive currents. (G) Outward currents at +60 mV in SMCs from MAs of Control (n = 8), AKAP5^SMCKO^ (n = 6), Cav1^SMCKO^ (n = 7), Piezo1^SMCKO^ (n = 7), and TRPV4^SMCKO^ (n = 11) mice before and after administration of Yoda1 (10 μM) or paxilline (1 μM) (***P < 0.001 vs. basal; ^##^P < 0.01 ^###^P < 0.001 vs. Control; 2-way ANOVA). (H) *Left*, representative PLA images showing SMC nuclei (blue) and Piezo1_SMC_:TRPV4_SMC_ colocalization (white puncta) in *en face* preparations of MAs from Cav1^fl/fl^, Cav1^SMCKO^, and AKAP5^SMCKO^ mice. *Right*, quantification of Piezo1_SMC_:TRPV4_SMC_ colocalization in MAs from Cav1^fl/fl^ (n = 6), Cav1^SMCKO^ (n = 6), and AKAP5^SMCKO^ (n = 6) mice (*P < 0.05 vs. Cav1^fl/fl^; 1-way ANOVA). (I) Cav1_SMC_-dependent mechanosensitive Piezo1_SMC_–TRPV4_SMC_–BK signaling promotes vasodilation and lowers blood pressure.

Pressure-induced (myogenic) constriction was higher in MAs from Piezo1^SMCKO^ mice than in MAs from control mice (Figure 4C), demonstrating a dilator effect of Piezo1_SMC_ channels under normal conditions. Intraluminal pressure increased the activity of TRPV4_SMC_ sparklets; this effect was absent in MAs from Piezo1^SMCKO^ mice (Figure 4D), suggesting that Piezo1_SMC_ channels mediate intraluminal pressure-induced activation of TRPV4_SMC_ channels. The Piezo1 agonist Yoda1 also activated outward currents through TRPV4_SMC_ channels in control mice but not in Piezo1^SMCKO^ mice (Figure 4E). Yoda1-activated BK currents were also reduced in SMCs from TRPV4^SMCKO^ and Piezo1^SMCKO^ mice (Figure 4F, G), further supporting Piezo1_SMC_–TRPV4_SMC_–BK channel signaling at dilator nanodomains.

Importantly, Piezo1_SMC_-activated TRPV4_SMC_ and BK currents were absent in Cav1^SMCKO^ mice (Figure 4E–G), further suggesting that Cav1 plays a critical role in promoting Piezo1_SMC_–TRPV4_SMC_–BK channel signaling at dilator nanodomains. Additionally, Piezo1_SMC_, TRPV4_SMC_, and BK channel currents elicited by direct channel agonists were not altered in Cav1^SMCKO^ mice (Figure 2E, 4B), suggesting that Cav1_SMC_ does not alter the number of functional Piezo1_SMC_, TRPV4_SMC_, or BK channels at the cell membrane. Overall, these data provide evidence for intraluminal pressure-activated Piezo1_SMC_–TRPV4_SMC_–BK channel signaling, which reduces myogenic constriction of MAs (Figure 4I).

A proximity analysis using PLA showed that Piezo1_SMC_ and TRPV4_SMC_ channels colocalize, and that Piezo1_SMC_:TRPV4_SMC_ colocalization is reduced in Cav1^SMCKO^ mice but not in AKAP5^SMCKO^ mice (Figure 4H). Moreover, α1AR-mediated vasoconstriction and activation of TRPV4_SMC_ sparklets were not altered in Piezo1^SMCKO^ mice (Figure S11), suggesting that Piezo1_SMC_ channels do not participate in α1AR-mediated constrictor signaling. In addition, Piezo1_SMC_ channel-activated BK currents were not altered in AKAP5^SMCKO^ mice. Together, these data provide evidence for dilator Piezo1_SMC_–TRPV4_SMC_–BK signaling facilitated by Cav1 and the absence of a role for Piezo1_SMC_ channels at AKAP5-dependent constrictor α1AR–TRPV4_SMC_ signaling nanodomains.

### Hyperactive AKAP5_SMC_:TRPV4_SMC_ nanodomains and impaired Cav1_SMC_:TRPV4_SMC_ nanodomains drive vasoconstriction and hypertension

Previous studies showed that hyperactive constrictor α1AR–TRPV4_SMC_ signaling and reduced TRPV4_SMC_–BK signaling contributes to vasoconstriction and blood pressure elevation in hypertension (*3*). We first tested the contribution of AKAP5_SMC_-dependent constrictor TRPV4_SMC_ signaling to pathogenesis of hypertension. Mean arterial pressure, recorded after 7 and 14 days of angiotensin II (Ang II, 1 μg/kg/min) infusion, was reduced in AKAP5^SMCKO^ mice when compared with control mice (Figure 5A), supporting a key role for AKAP5_SMC_ in Ang II-induced hypertension. Moreover, Ang II infusion-induced increase in blood pressure was lower in AKAP5^SMCKO^ mice than in control mice (Figure 5A), suggesting that a lower blood pressure in Ang II–infused AKAP5^SMCKO^ mice was not simply due to a lower starting blood pressure on day 0. Ang II infusion did not result in a significant change in heart rate in either group (Figure 5A). Additionally, acute treatment with α1AR antagonist prazosin (1 mg/kg intraperitoneally) (Figure 5B) or TRPV4 inhibitor GSK219 (1 mg/kg intraperitoneally) (Figure 5C) lowered blood pressure in control Ang II mice, and prazosin-or GSK219-induced decrease in blood pressure was reduced in Ang II AKAP5^SMCKO^ mice. Together, these data suggested that AKAP5_SMC_ is crucial for α1AR–TRPV4_SMC_-induced increase in blood pressure in hypertension.

**Figure 5.**
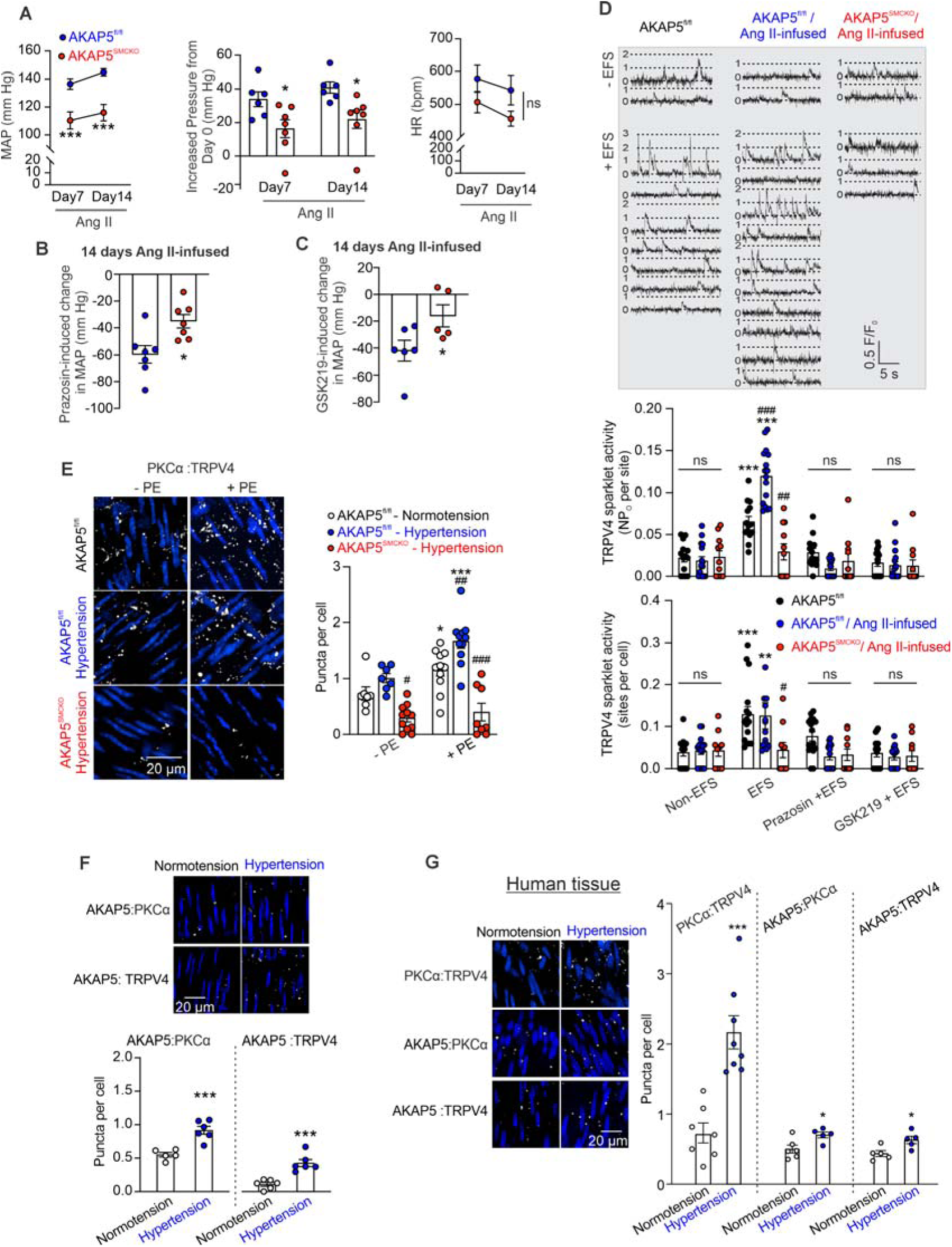
Enhanced signaling at AKAP5_SM_-dependent α1AR–PKCα–TRPV4_SMC_ nanodomains elevates blood pressure in hypertension. (A) *Left*, mean arterial pressure (MAP) in AKAP5^fl/fl^ (n = 6) and AKAP5^SMCKO^ (n = 6) mice, recorded before (day 0) and 7 and 14 days after Ang II infusion with osmotic minipumps (***P < 0.001 vs. AKAP5^fl/fl^; 2-way ANOVA). *Middle*, increased MAP in AKAP5^fl/fl^ (n = 6) and AKAP5^SMCKO^ (n = 6) mice after 7 and 14 days of Ang II infusion (*P < 0.05 vs. AKAP5^fl/fl^; 2-way ANOVA). *Right*, heart rate (HR) in AKAP5^fl/fl^ (n = 6) and AKAP5^SMCKO^ (n = 6) mice, analyzed before (day 0) and after 7 and 14 days of Ang II infusion (ns, not significant; 2-way ANOVA). (B) Change in MAP after a single bolus injection of prazosin (α1AR inhibitor, 1 mg/kg, ip) in 14-day Ang II mice (AKAP5^fl/fl^, n = 7; AKAP5^SMCKO^, n = 7) (*P < 0.05 vs. AKAP5^fl/fl^; unpaired t-test). (C) Changes in MAP after a single bolus injection of the TRPV4 inhibitor GSK219 (1 mg/kg, ip) in 14-day Ang II mice (AKAP5^fl/fl^, n = 6; AKAP5^SMCKO^, n = 5) (*P < 0.05 vs. AKAP5^fl/fl^; unpaired t-test). (D) Fluo-4AM–loaded, pressurized MAs were pretreated with CPA (20 μM) to eliminate interference from intracellular Ca^2+^ release, capsaicin (1 μM) to desensitize sensory nerves, atropine (10 μM) to block muscarinic receptors, and nifedipine (1 μM) to block L-type Ca^2+^ channels, before and after EFS, administered as 5-second trains of electrical pulses (50–150 V, 0.25-ms, 15 Hz). *Upper*, representative fractional fluorescence (F/F_0_) traces reflecting TRPV4_SMC_ sparklet activity in MAs from AKAP5^fl/fl^, Ang II AKAP5^fl/fl^, and Ang II AKAP5^SMCKO^ mice before and after EFS in the absence or presence of prazosin (1 μM; α1AR antagonist) or GSK219 (100 nM; TRPV4 antagonist). *Middle*, averaged TRPV4_SMC_ sparklet activity per site (NP_O_). *Lower*, averaged TRPV4_SMC_ sparklet sites per cell under basal conditions (CPA/capsaicin/atropine/nifedipine) in the absence or presence of prazosin (1 μM) or GSK219 (100 nM) in pressurized MAs from AKAP5^fl/fl^ (n = 14), Ang II AKAP5^fl/fl^ (n = 16), and Ang II AKAP5^SMCKO^ (n = 11) mice (**P < 0.01, ***P < 0.001 vs. Non-EFS; #P < 0.05, ^##^P < 0.01, ^###^P < 0.001 vs. AKAP5^fl/fl^; 2-way ANOVA). (E) *Left*, representative PLA images showing SMC nuclei (blue) and PKCα:TRPV4_SMC_ colocalization (white puncta) in *en face* preparations of MAs. *Right*, quantification of PKCα:TRPV4_SMC_ colocalization in MAs from control (normotensive) AKAP5^fl/fl^ mice and Ang II-infused (hypertensive) AKAP5^fl/fl^ and AKAP5^SMCKO^ mice before (AKAP5^fl/fl^, n = 11; Ang II AKAP5^fl/fl^, n = 11;Ang II AKAP5^SMCKO^, n = 8) and after (AKAP5^fl/fl^, n = 7; Ang II AKAP5^fl/fl^, n = 7; Ang II AKAP5^SMCKO^, n = 11) PE (1 μM) treatment (*P < 0.05, ***P < 0.001 vs. without PE; ^#^P < 0.05, ^##^P < 0.01, ^###^P < 0.001 vs. AKAP5^fl/fl^ ; 2-way ANOVA). (F) *Upper*, representative *in situ* PLA images showing SMC nuclei (blue), and AKAP5_SMC_:PKCα and AKAP5_SMC_:TRPV4_SMC_ colocalization in *en face* MAs from saline control and Ang II mice. *Lower*, quantification of AKAP5_SMC_:PKCα and AKAP5_SMC_:TRPV4_SMC_ colocalization in MAs from Ang II (n = 6) and control (n = 6) mice (***P < 0.001 vs. Normotension; unpaired t-test). (G) *Left*, representative *in situ* PLA images showing PKCα:TRPV4_SMC_, AKAP5_SMC_:PKCα, and AKAP5_SMC_:TRPV4_SMC_ colocalization (white puncta) in *en face* preparations of skeletal muscle arteries from normotensive and hypertensive individuals. *Right*, quantification of PKCα:TRPV4_SMC_ (Normotension, n = 7; Hypertension, n = 8), AKAP5_SMC_:PKCα (Normotension, n = 6; Hypertension, n = 5) and AKAP5_SMC_:TRPV4_SMC_ (Normotension, n = 6; Hypertension, n = 5) colocalization in arteries from normotensive and hypertensive individuals (*P < 0.05 vs. Normotension; unpaired t-test).

Sympathetic nerve stimulation-induced TRPV4_SMC_ sparklet activity was higher in Ang II mice than that in saline-infused control mice. Moreover, nerve stimulation-induced TRPV4_SMC_ sparklet activity in Ang II mice was abolished by inhibitors of α1AR and TRPV4 (prazosin and GSK219, respectively), suggesting increased activation of α1AR–TRPV4_SMC_ signaling by sympathetic stimulation in hypertension. Notably, nerve stimulation was unable to increase TRPV4_SMC_ sparklet activity in Ang II-infused AKAP5^SMCKO^ mice (Figure 5D), providing evidence that AKAP5_SMC_ is critical for increased α1AR–TRPV4_SMC_ constrictor signaling in hypertension.

PLA studies showed increased PKCα:TRPV4_SMC_ colocalization in arteries from control Ang II mice. Moreover, α1AR agonist phenylephrine (1 μM) increased PKCα:TRPV4_SMC_ colocalization in control Ang II mice but not in AKAP5^SMCKO^ Ang II mice (Figure 5E). AKAP5_SMC_:PKCα and AKAP5_SMC_:TRPV4_SMC_ colocalization was also increased in arteries from Ang II mice (Figure 5F) compared to control mice. PLA studies in human samples also showed an increase in PKCα:TRPV4_SMC_, AKAP5_SMC_:PKCα and AKAP5_SMC_:TRPV4_SMC_ colocalization in hypertensive human subjects compared to normotensive individuals (Figure 5G). Collectively, these data suggested increased AKAP5_SMC_-dependent α1AR-PKCα-TRPV4_SMC_ signaling in hypertension.

Notably, mean arterial pressure, measured after 7 and 14 days of Ang II infusion, and Ang II infusion-induced increase in blood pressure were not significantly different between Cav1_SMCKO_ mice than in control mice (Figure 6A). These data are consistent with previous findings that dilator TRPV4_SMC_–BK signaling is absent in hypertension (*3*), and therefore, SMC-specific deletion of Cav1 may not produce additional effects on blood pressure. However, acute administration of the TRPV4 inhibitor GSK219 (1 mg/kg, intraperitoneally) (Figure 6B) produced a greater reduction in blood pressure in Ang II-infused Cav1_SMCKO_ mice than in Ang II-infused control mice. This effect could be attributed to the selective inhibition of hyperactive constrictor TRPV4_SMC_ signaling in absence of the dilator TRPV4_SMC_ signaling in Cav1_SMCKO_ mice.

**Figure 6.**
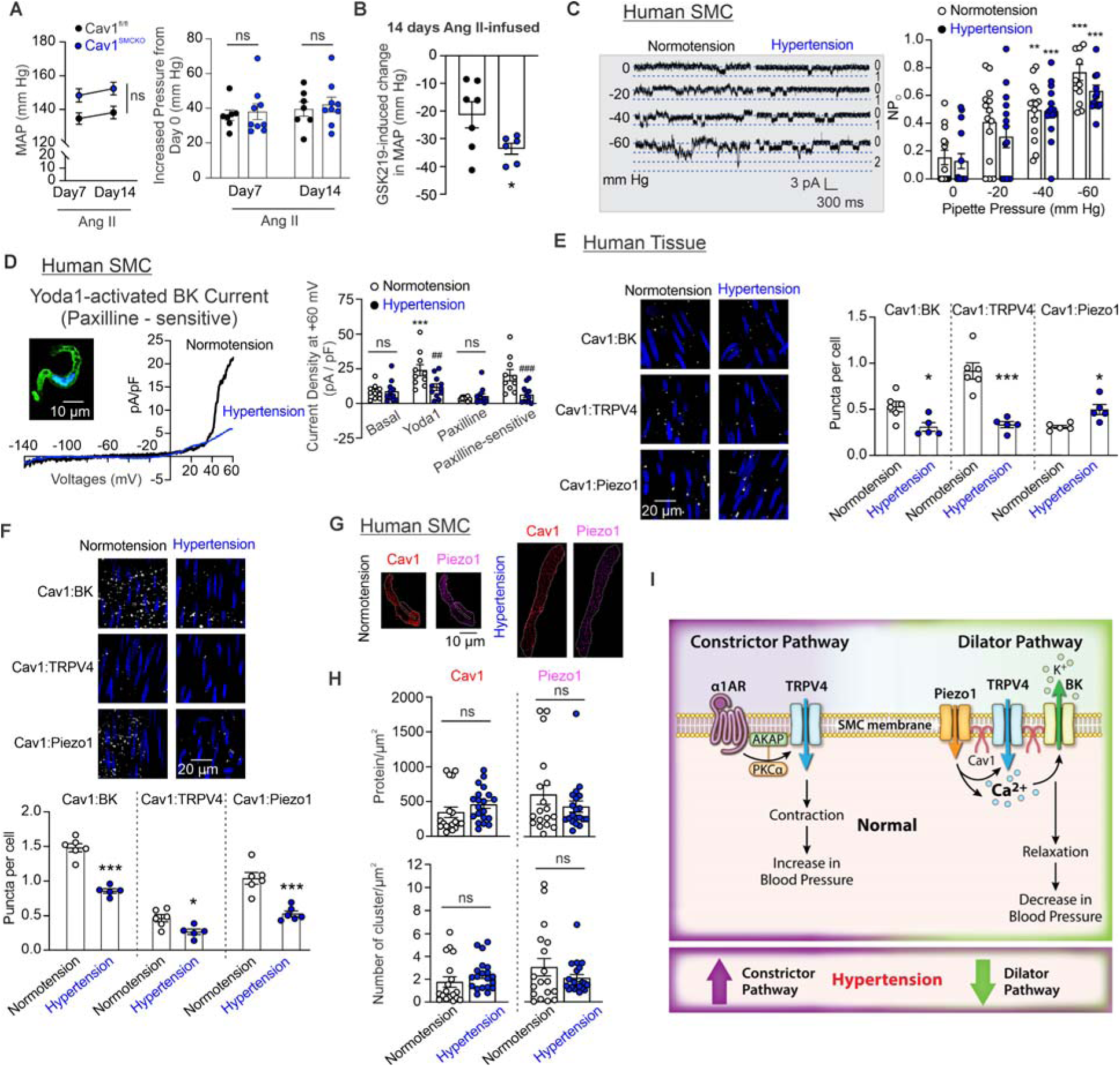
Cav1–dependent dilator Piezo1_SMC_–TRPV4_SMC_–BK signaling is impaired in hypertensive individuals and Ang II mice. (A) *Left*, mean arterial pressure (MAP) in Cav1^fl/fl^ (n=7) and Cav1^SMCKO^ (n=9) mice, recorded before the start of Ang II infusion (day 0) and 7 and 14 days after Ang II infusion through osmotic minipumps (ns, not significant; 2-way analysis of variance). *Right*, increased MAP in Cav1^fl/fl^ (n=7) and Cav1^SMCKO^ (n=9) mice after 7 and 14 days of Ang II infusion (ns, not significant; 2-way analysis of variance). (B) Change in MAP after a single bolus injection of GSK219 (1 mg/kg intraperitoneally) in 14 days of Ang II infusion (Cav1^fl/fl^, n=7; Cav1^SMCKO^, n=6) (*P<0.05 vs Cav1^fl/fl^; unpaired t-test). (C) *Left*, representative traces showing mechanically activated Piezo1 channel currents in freshly isolated SMCs from normotensive and hypertensive individuals. Gradually increasing suction (0, -20, -40, and -60 mm Hg) elicited inward currents through Piezo1 channels. Dotted lines represent quantal levels (single-channel amplitudes) determined from all-points histograms. *Right*, Piezo1 channel activity (NP_O_, where N is the number of channels and P_O_ is the open state probability) with gradually increased suction in SMCs from normotensive (n = 14) and hypertensive (n = 15) individuals (**P < 0.01, ***P < 0.001 vs. 0 mm Hg; ns, not significant; 2-way ANOVA). (D) *Inset,* widefield image of a SMC from a human skeletal muscle artery, stained for α-actin (green) and counterstained with the nuclear dye DAPI (blue). The fusiform shaped SMCs were used in patch-clamp experiments. *Left*, representative traces of BK channel currents induced by the Piezo1 agonist Yoda1 (10 μM), measured as paxilline-sensitive currents in SMCs from human subjects (n = 1–2 SMCs per subject). *Right*, outward currents at +60 mV in SMCs from normotensive (n = 10) and hypertensive (n = 11) subjects before and after administration of Yoda1 (10 μM) or Yoda1+paxilline (1 μM) (***P < 0.001 vs. basal; ^##^P < 0.01 ^###^P < 0.001 vs. Normotension; ns, not significant; 2-way ANOVA). (E) *Left*, representative *in situ* PLA images showing Cav1_SMC_:BK, Cav1_SMC_:TPRV4_SMC_, and Cav1_SMC_: Piezo1_SMC_ colocalization (white puncta) in *en face* preparations of skeletal muscle arteries from normotensive (control) and hypertensive individuals. *Right*, quantification of Cav1_SMC_:BK, Cav1_SMC_:TPRV4_SMC_, and Cav1_SMC_: Piezo1_SMC_ colocalization in arteries from normotensive (control; n = 6) and hypertensive (n = 5) individuals (*P < 0.05, ***P < 0.001 vs. Normotension; unpaired t-test). (F) *Upper*, representative *in situ* PLA images showing Cav1_SMC_:BK, Cav1_SMC_:TPRV4_SMC_, and Cav1_SMC_: Piezo1_SMC_ colocalization (white puncta) in *en face* preparations of MAs from Ang II-infused and saline-treated (control) mice. *Lower*, quantification of Cav1_SMC_:BK, Cav1_SMC_:TPRV4_SMC_, and Cav1_SMC_: Piezo1_SMC_ colocalization in MAs from Ang II (n = 6) and control (n = 5) mice (*P < 0.05, ***P < 0.001 vs. Normotension; unpaired t-test). (G) Representative TIRF-SMLM images acquired using spectral demixing showing localization maps for Cav1 and Piezo1 channels at the SMC plasma membrane in SMCs from normotensive and hypertensive individuals. (H) Summary of TIRF-SMLM localization maps showing molecular density (*upper*) and cluster density (*lower*) of Cav1 and Piezo1 channels at the plasma membrane in SMCs from normotensive (n = 18) and hypertensive (n = 20) individuals (1–2 SMCs/participant; unpaired t-test). (I) Schematic diagram showing AKAP5_SMC_-dependent constrictor TRPV4_SMC_ nanodomains and Cav1_SMC_-based dilator TRPV4_SMC_ nanodomains, and their imbalance in hypertension.

Studies of Cav1_SMC_-based dilator nanodomains in skeletal muscle arteries from human subjects showed that stretch-induced activation of Piezo1_SMC_ channels is not different between SMCs from normotensive or hypertensive subjects, indicating that the number of functional channels at the cell membrane is not altered in hypertension (Figure 6C). However, Piezo1-activated BK currents (paxilline-sensitive currents) were reduced in hypertensive patients compared to normotensive individuals (Figure 6D). PLA studies showed that Cav1_SMC_:BK, Cav1_SMC_:TRPV4_SMC_, and Cav1_SMC_:Piezo1_SMC_ colocalization was impaired in SMCs from hypertensive subjects (Figure 6E) and hypertensive mice (Figure 6F) compared to respective normotensive controls. However, TIRF-SMLM images suggested that the molecular density and cluster density of Piezo1_SMC_ and Cav1_SMC_ is not affected in hypertensive subjects (Figure 6G-H; negative control in Figure S12), indicating disrupted colocalization rather than reduced membrane expression as a mechanism for impaired Cav1_SMC_-dependent Piezo1_SMC_–TRPV4_SMC_–BK signaling in hypertension (Figure 6I).

## Discussion

Our results identify the cellular mechanisms underlying the formation of spatially separated TRPV4_SMC_ signaling nanodomains and explain their opposite effects on blood pressure. These findings support the broader concept that signaling through specific partners in the nano-scale neighborhood enables an ion channel to produce divergent physiological effects. At the dilator nanodomain, Cav1 promotes the formation of a mechanosensitive Piezo1_SMC_:TRPV4_SMC_:BK channel signaling module that mediates vasodilation and lowers blood pressure (Figure 6G). In contrast, the constrictor nanodomain relies on AKAP5_SMC_-dependent PKCα– TRPV4_SMC_ signaling to mediate sympathetic vasoconstriction and elevate blood pressure. In hypertensive human subjects and a mouse model of hypertension, AKAP5_SMC_-dependent nanodomains are hyperactive, whereas Cav1-based nanodomains are hypoactive. This imbalance between constrictor and dilator nanodomains amplifies vasoconstriction and drives blood pressure elevation in hypertension (Figure 6I). Moreover, impaired nanoscale colocalization of proteins with AKAP5_SMC_ or Cav1_SMC_ underlies this imbalance, providing a mechanistic basis for increased vasoconstriction and blood pressure in hypertension.

L-type Ca^2+^ channels (LTCC), which are well-known for their role in SMC contraction, are upregulated in hypertension (*25–27*). Previous studies demonstrated that AKAP5_SMC_-anchored PKC also increases LTCC activity (*14, 28*), raising the possibility of a functional overlap with constrictor TRPV4_SMC_ channels. However, our published data indicate that LTCC inhibition does not affect TRPV4_SMC_ channel activity (*3*), supporting the concept that these channels operate independently. Collectively, our findings indicate that AKAP5_SMC_–PKC– LTCC signaling may be spatially distinct from AKAP5_SMC_–PKC–TRPV4_SMC_ signaling and may be activated by different physiological stimuli: whereas AKAP5_SMC_–PKC–LTCC signaling has been associated with pressure-induced constriction (*14*), AKAP5_SMC_–PKC–LTCC signaling mediates sympathetic vasoconstriction.

Sympathetic stimulation has also been shown to trigger ATP efflux through pannexin 1, promoting SMC contraction (*29, 30*). These observations raise the intriguing possibility that pannexin 1 may be upstream of PKCα–TRPV4_SMC_ signaling. If ATP serves to promote constrictor PKCα–TRPV4_SMC_ signaling, it could represent a novel mechanism for sympathetic regulation of SMC Ca^2+^ signaling and vascular resistance. Additionally, previous studies showed that SMC-specific Cav1 knockout reduces α1AR-mediated constriction in thoracodorsal arteries (*31*). The apparent discrepancy in the role of Cav1_SMC_ in regulating α1AR-mediated vasoconstriction could be explained by the use of different experimental conditions, including the size of the artery/vascular bed (resistance-sized mesenteric arteries with myogenic constriction in the current study versus pre-constricted thoracodorsal arteries in (*31*)) and tamoxifen administration (intraperitoneal injection, which may affect mesenteric vasculature, and therefore, blood pressure, versus dietary tamoxifen used in the current study.

Our findings provide evidence that mechanosensitive Piezo1 channels oppose pressure-induced vasoconstriction under physiological conditions, revealing a new vasodilator mechanism in SMCs. The precise mechanism by which Piezo1 channels activate TRPV4 channels in SMCs and other cell types is not clear (*32, 33*). Studies in endothelial cells indicated that Piezo1-activation of TRPV4 channels could be occurring through activation of phospholipase A2 (PLA2) (*34*), a possibility that has not been investigated in SMCs. Previous studies using non-inducible Piezo1^SMCKO^ mice ruled out a role for Piezo1 channels in regulating pressure-induced vasoconstriction (*35*). In contrast, our studies using inducible Piezo1^SMCKO^ mice show that Piezo1 channels are a critical component of the Cav1-based dilator nanodomain, which limits myogenic constriction. Cav1 is known to organize signaling components within specialized membrane microdomains called caveolae, suggesting that caveolar localization of the dilator nanodomain may be crucial for its function. However, whether caveolae serve as the primary microenvironment for the dilator Piezo1_SMC_*–*TRPV4_SMC_–BK nanodomain remains unknown. Emerging evidence suggests that Cav1 outside caveolae can integrate mechanosensitive signaling (*36*). Moreover, mechanical stress rapidly induces caveolar disassembly, releasing Cav1 scaffolds that exhibit enhanced membrane diffusion and interactions with signaling proteins (*37*). Thus, it is plausible that pressure-induced redistribution of Cav1 scaffolds promotes the formation of dilator Piezo1_SMC_*–* TRPV4_SMC_–BK nanodomains.

Our data show that Cav1-dependent dilator Piezo1_SMC_*–*TRPV4_SMC_–BK channel signaling is impaired in hypertension. Notably, the amount of Cav1, Piezo1, TRPV4, and BK channels at the SMC membrane remained unchanged, implicating mis-localization of these channels relative to Cav1 as a key mechanism underlying nanodomain dysfunction. The causative mechanisms for enhanced AKAP5*–*PKCα*–*TRPV4_SMC_ signaling and reduced Cav1-dependent Piezo1_SMC_*–*TRPV4_SMC_–BK channel signaling are not known. Recent studies have shown that AKAP5_SMC_ expression is increased in hypertensive rats (*38*). Increased AKAP5_SMC_ expression could lead to enhanced constrictor TRPV4_SMC_ signaling, although this possibility has not been verified. Moreover, alterations in membrane lipid composition or caveolar integrity could contribute to diminished dilatory signaling and elevated blood pressure in hypertensive states. Whether hypertension disrupts caveolar morphology, modifies Cav1 conformation, or impairs its scaffolding capacity for mechanosensitive channels remains an important question for future investigation. Elucidating how Cav1-dependent scaffolding is altered in hypertension may uncover novel mechanisms of impaired mechanotransduction and identify therapeutic strategies to restore physiological dilatory signaling.

Although a physiological role for Piezo1_SMC_ channels has not been previously described, these channels have been implicated in vascular remodeling and elevated arterial pressure in systemic and pulmonary hypertension (*35, 39*). Moreover, total expression of Piezo1_SMC_ channels is increased in pulmonary hypertension (*39*). In contrast, our data demonstrate that the number of Piezo1 molecules at the SMC plasma membrane remains unchanged in hypertension, while pressure-induced dilatory Piezo1 activity is markedly reduced. Thus, it is conceivable that the mis-localization of Piezo1 channels from Cav1-based dilator nanodomains drives the functional transition of these domains from dilator to constrictor signaling. Whether Piezo1 channels participate in additional nanodomains that promote vascular remodeling in hypertension is not known.

Additional intracellular proteins likely contribute to the functional specificity of distinct signaling nanodomains in SMCs. For example, ryanodine receptors (RyR) at the sarcoplasmic reticulum (SR) membrane colocalize with BK channels and promote vasodilation through Ca^2+^ release and subsequent BK channel activation (*9*). Given this well-established spatial coupling, it is plausible that RyRs act as downstream targets of Piezo1_SMC_–TRPV4_SMC_ signaling, coupling mechanosensation with BK channel activation. Indeed, previous studies reported reduced RyR Ca^2+^ signaling activity in TRPV4^SMCKO^ mice (*3*). Conversely, Ca^2+^ release through IP_3_ receptors (IP_3_Rs) at the SR membrane plays a key role in SMC contraction. Prior work showed that IP_3_R Ca^2+^ signals are also diminished in TRPV4^SMCKO^ mice (*3*), suggesting that RyR and IP_3_R may serve as downstream effectors of dilator and constrictor nanodomains, respectively, further contributing to their functional specificity. Furthermore, differential regulation of myosin light chain kinase (MLCK) by these spatially distinct nanodomains, and its impact on SMC contraction, cannot be excluded.

In summary, this study explains the functionally opposite effects of distinct subpopulations of TRPV4 channels at the SMC membrane by identifying a vasodilator nanodomain containing Cav1-based Piezo1_SMC_– TRPV4_SMC_–BK signaling and a constrictor nanodomain involving AKAP5_SMC_-dependent α1AR–PKCα– TRPV4_SMC_ signaling. The spatial separation of these opposing nanodomains at the SMC plasma membrane provides a fundamental mechanism for fine-tuning vascular tone and blood pressure. In hypertension, altered mis-localization of the signaling elements at the two nanodomains leads to hyperactivation of constrictor nanodomains and impairment of dilator nanodomains, shifting the balance towards vasoconstriction and elevating blood pressure. These findings advance our understanding of how distinct scaffolding proteins promote nanoscale organization of TRPV4 channels with their signaling partners to fine-tune vascular function and reveal new mechanistic targets for restoring physiological signaling in hypertensive states.

## Materials and Methods

### Animal protocols

Animal care was performed in accordance with institutional guidelines and animal protocols approved by the University of Virginia Animal Care and Use Committee (protocols 4100 and 4120). Food and water were provided *ad libitum* until the study was completed. C57BL6/J mice (Jackson Laboratory, Bar Harbor, ME, USA) or mice on a C57BL6/J background (10–14 weeks old) were fed a normal chow diet (#7912; Envigo, Indianapolis, IN, USA), weaned at 3 weeks of age, and kept under standard conditions (12-hour light/dark cycles and 21 ± 2°C). Both male and female mice were used in this study. For harvesting mesenteric arteries (MAs), mice were euthanized with pentobarbital (90 mg/kg, i.p.; Diamondback Drugs, UVA Hospital Pharmacy) followed by cervical dislocation.

For all *in vivo* experiments, random assignment of animals to groups was performed by an independent team member without knowledge of assigned treatments, and data were analyzed by an investigator blinded to group-and treatment-identifying information. All experiments were performed in at least two independent batches.

### Generation of knockout mice

*AKAP5^fl/fl^* (*40*)*, Cav1^fl/f^ ^l^*(*41*), *Piezo1^fl/^*^fl^ (*42*) and *Trpv4^fl/fl^* (*43*) and mice were crossed with *Myh11Cre^ERT2^RAD* (*44*) mice to generate tamoxifen-inducible, SMC-specific knockout (SMCKO) mice. SMC-specific knockout was induced in 6-week-old *AKAP5^fl/fl^ Myh11-*Cre^ERT2^RAD, *Cav1^fl/fl^ Myh11-*Cre^ERT2^RAD, *Piezo1^fl/^*^fl^ *Myh11-*Cre^ERT2^RAD, and *Trpv4^fl/fl^ Myh11-*Cre^ERT2^RAD mice by feeding tamoxifen (40 mg/kg/d; Envigo Diet TD.130856) (Table S1) for 14 days. Cre-negative tamoxifen-fed *AKAP5^fl/fl^*, *Cav1^fl/fl^*, *Piezo1^fl/^*^fl^ and *Trpv4^fl/fl^* mice were used as controls. All transgenic mice were backcrossed onto a C57BL/6 background for at least 10 generations. Mice were used for experiments after a 1-week tamoxifen-washout period followed by a normal chow diet. Cell-specific deletion was validated by immunofluorescence as described below.

### Genotyping

Genomic DNA was extracted from ear or tail tissue samples by treating with DirectPCR lysis reagents (200 mL for tail and 100 mL for ear samples) containing 0.2 mg/mL proteinase K at 55°C for 3 hours or until no tissue clumps were observed. The crude lysate was then incubated at 85°C for 45 minutes to inactivate the proteinase K. After centrifuging crude DNA samples at 10,000′ g for 2 minutes at room temperature, 0.5 mL of lysate was used for each 25-mL PCR reaction. PCR was performed on a T100 Thermal Cycler (Bio-Rad Laboratories) using 2x Platinum mix, 1 μM 5’ and 3’ primers, and ∼100–250 ng genomic DNA. Reaction products were resolved on a 2–3% agarose gel containing 0.2 µg/µL ethidium bromide in TAE buffer (40 mM Tris base, 20 mM acetic acid, 1 mM EDTA) at 90V. Gels were visualized using 302 nm UV light, and the sizes of PCR products were calibrated using a 100-bp DNA Ladder (New England BioLabs). Mice were genotyped using the primer pairs listed in Tables S2 and S3. All primers were obtained from Eurofins Genomics.

### Drugs and chemicals

Cyclopiazonic acid (CPA; SERCA inhibitor), GSK2193874 (GSK219; TRPV4 inhibitor), GSK1016790A (GSK101; TRPV4 agonist), and Fluo 4 AM (Ca^2+^ indicator) were purchased from Invitrogen. All chemicals and drugs are listed in Table S4.

### Radiotelemetric blood pressure measurement

Continuous blood pressure measurements were performed using Ponemah 6.42 software (Data Sciences International, St. Paul, MN, USA), as described previously (*3, 45*). Mice were anesthetized with isoflurane (1.5%), and a radiotelemetry catheter (HD-X10; Data Sciences International, St. Paul, MN, USA) was inserted in the left carotid artery. The radiotransmitter was placed in a subcutaneous pouch along the flank. After surgery, mice were allowed to recover for 7 days to regain normal circadian rhythms before initiating arterial pressure measurements. Following the recovery period, baseline systolic pressure, diastolic and mean arterial pressure, and heart rate were recorded continuously over 48 hours at 1-minute intervals. Baseline daytime and nighttime recordings were obtained by averaging values over 2 days (6 AM to 6 PM) and 2 nights (6 PM to 6 AM), respectively. *AKAP5*^fl/fl^, *AKAP5*^SMCKO^, *Cav1*^fl/fl^, and *Cav1*^SMCKO^ mice were given a single bolus intraperitoneal injection of the α1AR agonist phenylephrine (10 mg/kg); angiotensin II (Ang II)-infused *AKAP5*^fl/fl^, *AKAP5*^SMCKO^, *Cav1*^fl/fl^, and *Cav1*^SMCKO^ mice were given a single bolus intraperitoneal injection of the α1AR agonist phenylephrine (10 mg/kg), α1AR antagonist prazosin (1 mg/kg), or TRPV4 inhibitor GSK219 (1 mg/kg). Sterile saline solution (0.9%) or DMSO (dimethyl sulfoxide) was used as a vehicle; mice injected with vehicle only were used as controls group for statistical comparisons. Blood pressure and heart rate were recorded for 1 hour at 5-minute intervals. Blood pressures obtained 15 minutes after injection were compared between groups. Experiments were performed in a blinded manner.

### Isolation of SMCs from arteries

SMCs were freshly isolated from third-order MAs (∼ 100 μm) from mice or human skeletal muscle arteries. Briefly, artery segments were transferred to a 12 x 75 mm borosilicate glass culture tube containing 1 mL dissociation solution (145 mM NaCl, 4 mM KCl, 1 mM MgCl_2_, 10 mM HEPES, 0.05 mM CaCl_2_ 10 mM glucose) and 0.5 mg/mL bovine serum albumin (BSA) and then incubated for 10 minutes at room temperature (∼24°C). This solution was then replaced with 1 mL dissociation solution containing 1 mg/mL papain (MilliporeSigma) and 0.5 mg/mL dithiothreitol (MilliporeSigma) at 37°C for 6 minutes. Thereafter, 0.5 mL of the papain solution was carefully removed without displacing artery segments and replaced with 0.5 mL dissociation solution containing 2 mg/mL collagenase type IV (Worthington Biochemical Corporation, Lakewood, NJ, USA), 0.5 mg/ml elastase (MilliporeSigma), and 1 mg/ml soybean trypsin inhibitor (MilliporeSigma) and incubated at 37°C for 8 minutes. The enzyme solution was then removed and replaced with cold dissociation solution containing BSA. The tube containing digested arteries was placed on ice, and the solution was gently triturated every 15 minutes for 1 hour to yield a single-cell suspension.

### Whole-cell patch-clamp electrophysiology

Patch electrodes were pulled with a Narishige PC-100 puller (Narishige International USA, Inc, Amityville, NY) and polished using a MicroForge MF-830 polisher (Narishige International USA). The pipette resistance was (3–5 ΩM). Data were acquired using a Multiclamp 700B amplifier connected to a Digidata 1550B system and analyzed using Clampfit 11.1 software (Molecular Devices, San Jose, CA, USA) and MATLAB R2018a (MathWorks, Natick, MA, USA). TRPV4 channel currents were recorded from freshly isolated SMCs as described previously (3). Outward currents through TRPV4 channels induced by phenylephrine (1 μM) or Yoda1 (10 μM) were assessed using the whole-cell configuration of the patch-clamp technique in the presence of ruthenium red, included to prevent Ca^2+^ influx through TRPV4 channels and activation of BK channels. The intracellular solution consisted of 20 mM CsCl, 100 mM Cs aspartate, 1 mM MgCl_2_, 4 mM ATP, 0.08 mM CaCl_2_, 10 mM BAPTA, and 10 mM HEPES, pH 7.2 (adjusted with CsOH). TRPV4 currents were measured using a voltage-clamp protocol in which voltage-ramp pulses (−100 mV to +100 mV) were applied over 200 ms from a holding potential of -50 mV. Currents were measured before and 5 minutes after treatment with GSK219 (100 nmol/L), with GSK219-sensitive currents being indicative of TRPV4 channel currents. Phenylephrine (1 μM)-induced outward currents through TRPV4 channels were also assessed in the absence/presence of the PKCα/β inhibitor Gö 6976 (1 μM). Maximum TRPV4 channel activity was assessed by applying 100 nM GSK101. Activation of BK channels by TRPV4 or Piezo1 channel signaling was evaluated by applying GSK101 (30 nM) or Yoda1 (10 μM), respectively. BK currents were measured using a voltage ramp from -140 mV to +60 mV over 250 ms. Currents were measured before and 5 minutes after treatment with paxilline (1 μM), with paxilline-sensitive currents corresponding to BK currents. The BK channel activator GoSlo SR 5-69 (1 μM) was also used to assess direct activation of BK channels.

### Isolation of human skeletal muscle tissue from hypertensive and non-hypertensive individuals

Skeletal muscle tissue was obtained from hypertensive and non-hypertensive subjects during spinal surgeries (Table S5), as approved by the University of Virginia Institutional Review Board (Protocol #18699). Informed consent was obtained from each subject as per the protocol. Hypertensive patients were designated as those with a documented pre-operative history of treatment with antihypertensive agents. Patient histories were reviewed to ensure that the medication was not started for other indications (e.g., low-dose ACE inhibitor for diabetic nephropathy, initiated in one patient). In addition, patients in the “non-hypertensive” group were confirmed not to be overtly hypertensive based on pre-operative blood pressure measurements. For this group, borderline measurements with systolic blood pressures in the 140 mm Hg range were still accepted as “non-hypertensive.” Individuals treated with α1AR antagonists as antihypertensive medication were excluded from the study. Detailed clinical information, including antihypertensive medications, reason for surgery and other medication, is presented in Table S5. Arteries were dissected from muscle tissue, placed in cold HEPES-PSS, and then used for *in situ* proximity ligation assay (PLA) or enzymatic isolation of SMCs for whole-cell patch-clamp electrophysiology.

### Pressure myography

Third-order MAs (∼100 µm) were cannulated onto two glass micropipettes on a custom-made pressure myography chamber (Instrumentation and Model Facility, University of Vermont, Burlington, VT, USA), pressurized to 80 mmHg with a pressure servo controller peristaltic pump (Living Systems Instrumentation, St Albans, VT, USA), and superfused at 37°C with PSS (119 mM NaCl, 4.7 mM KCl, 1.2 mM KH_2_PO_4_, 1.2 mM MgCl_2_ 160 hexahydrate, 2.5 mM CaCl_2_ dihydrate, 7 mM dextrose, and 24 mM NaHCO_3_) maintained at pH 7.4 by bubbling with 21% O_2_ and 5% CO_2_. MAs were allowed to develop spontaneous pressure-induced constriction (myogenic tone) at 80 mmHg before initiating experimental treatments. Dilation to NS309 (1 μM), a direct opener of intermediate/small-conductance Ca^2+^-sensitive K^+^ (IK/SK) channels, was used to functionally assess the health of endothelial cells. Changes in diameter in response to cumulative administration of phenylephrine (10 nM to 10 μM) or GSK101 (3, 10, 30, 100 nM), were recorded. Some groups of MAs were pretreated with GSK219 (100 nM) or paxilline (1 μM) for 10 minutes before cumulative exposure to phenylephrine or GSK101. MAs were exposed to each concentration of agonist until a stable diameter was attained (5–10 minutes). At the end of each experiment, maximum passive diameter was evaluated by incubating MAs with Ca^2+^-free PSS (119 mM NaCl, 4.7 mM KCl,1.2 mM KH_2_PO_4_,1.2 mM MgCl_2_, 7 mM glucose, 24 mM NaHCO_3_, 5 mM EGTA; pH 7.4). Internal diameter was recorded at 10 frames/s using a CCD camera and analyzed using IonOptix edge-detection software (IonOptix LLC, Westwood, MA, USA). Changes in diameter in response to agonists and antagonists were normalized to the resting diameter and expressed as a percentage.

Myogenic constriction at a perfusion pressure of 80 mm Hg was calculated as:

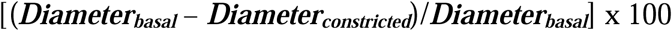

Myogenic constriction at perfusion pressures of 20, 40, 60, 80, 100, and 120 mm Hg was calculated as:

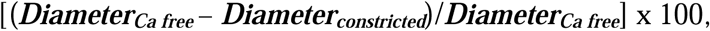

where ***Diameter_constricted_***is the diameter of the artery after drug-induced constriction or pressure-induced myogenic constriction, and ***Diameter_Ca free_***is the maximum passive diameter at the relative perfusion pressure.

### Endothelium-denuded artery for pressure myography

One side of a third-order MA (∼100 µm) was cannulated onto s glass micropipette on a custom-made pressure myography chamber. Air bubbles were passed through the artery for 50 seconds. The other side of the MA was then cannulated and the pressure myography protocol was performed as described above.

### Electric field stimulation (EFS)

Two platinum electrodes (0.25-mm diameter, World Precision Instruments) were placed in parallel on either side of a MA and connected to a Grass S88 stimulator. After equilibration at 37°C, perivascular nerves of pressurized (80 mmHg) arteries were stimulated by trains of electrical pulses (50– 150 V, 0.25-ms pulse, 15 Hz, 5-second train duration) (*46, 47*). A train duration of 5 seconds was chosen to minimize tissue movement for Ca^2+^ measurements. CPA (20 μM), capsaicin (1 μM; sensory nerve desensitizer), atropine (10 μM; muscarinic receptor antagonist), and nifedipine (1 μM; L-type Ca^2+^ channel antagonist) were prepared as stock solutions and diluted in the superfusate.

### Ca^2+^ imaging

Ca^2+^-imaging studies were performed as described previously (*3*). Briefly, third-order MAs from mice (∼100 µm) were incubated with Fluo-4 AM (10 μM) and pluronic acid (0.04%) at 37°C for 1 hour, after which they were cannulated and pressurized to 80 mmHg. Ca^2+^ images were acquired at 30 frames per second using an Andor Revolution WD (with Borealis) spinning-disk confocal imaging system (Andor Technology) comprising an upright Nikon microscope with a 40X water-dipping objective (numerical aperture, 0.8) and an electron-multiplying CCD camera. MAs were superfused with PSS (pH 7.4), and all experiments were performed at 37°C. Fluo-4 was excited using a 488 nm solid-state laser, and emitted fluorescence was captured using a 525/36 nm band-pass filter. Intracellular Ca^2+^-release signals and extracellular Ca^2+^-influx signals were eliminated by treating MAs with the sarco-endoplasmic reticulum Ca^2+^-ATPase (SERCA) inhibitor CPA (20 μM) and L-type Ca^2+^ channel inhibitor nifedipine (1 μM), respectively, for 10 minutes at 37°C prior to imaging. (We previously showed that CPA per se does not alter the activity of TRPV4 sparklets in endothelial cells (*48*).) For sympathetic nerve stimulation (EFS) described above, capsaicin (1 μM; sensory nerve desnsitizer) and atropine (10 μM; muscarinic receptors antagonist) were added to the incubation solution, and Ca^2+^ signals were recorded without/with EFS in the absence/presence of prazosin (1 μM; α1ARs antagonist) or GSK219 (100 nM). The influence of the pharmacological agent phenylephrine and intraluminal pressure (from 20 mmHg to 80 mmHg; 5 minutes at each pressure step) on TRPV4 sparklet activity was further evaluated. SMC TRPV4 sparklet inhibition was confirmed after a 10-minute incubation with GSK219 (100 nM). Ca^2+^ images were analyzed using custom-designed SparkAn software developed by Dr. Adrian Bonev (University of Vermont). Fractional fluorescence traces (F/F_0_) were obtained by placing a 1.7 μm^2^ (5 × 5 pixels) region of interest (ROI) at the peak event amplitude. Representative F/F_0_ traces were filtered using a Gaussian filter and a cutoff corner frequency of 4 Hz.

### Analysis of the activity of SMC TRPV4 sparklets

Ca^2+^ signals were assessed as increased fluorescence relative to baseline fluorescence obtained by averaging 10 quiescent images prior to stimulation. TRPV4 Ca^2+^ sparklets were assessed based on previously established methods (*3, 19*). Average TRPV4 sparklet activity is defined as NP_O_, where N is the number of TRPV4 channels per site and P_O_ is the open state probability of the channel. NP_O_ was calculated using the Single Channel Search module of Clampfit, quantal amplitudes derived from all-points histograms (0.3 ΔF/F_0_ for Fluo-4-loaded MAs), and the following equation:

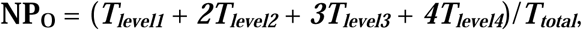

where T represents the dwell time at each quantal level and ***T_total_*** is the total recording duration. Average NP_O_ per site was obtained by averaging the NP_O_ for all sites in a field of view. NP_O_ per site from all fields in an artery were averaged to obtain NP_O_ per site for that artery. The total number of sites per cell was obtained by dividing the number of sparklet sites in a field by the number of cells in that field. Sparklet sites per cell for all fields were averaged to obtain sparklet sites per cell for that artery.

### Single-channel patch-clamp electrophysiology

Patch-clamp electrodes were pulled to a resistance of 4–6 MΩ with a Narishige PC-100 puller (Narishige International USA Inc.) and polished using a MicroForge MF-830 polisher (Narishige International USA, Inc.). Single-channel currents were recorded in the cell-attached configuration using a Multiclamp 700B amplifier connected to a Digidata 1550B system and analyzed using Clampfit 11.1 software (Molecular Devices) and MATLAB R2018a (MathWorks). The bath solution consisted of 10 mM HEPES, 134 mM NaCl, 6 mM KCl, 2 mM CaCl_2_, 10 mM glucose and 1 mM MgCl_2_ (adjusted to pH 7.4 with NaOH). The pipette solution consisted of 3 mM KCl, 134 mM NaCl, 1 mM MgCl_2_, 10 mM HEPES, 4 mM glucose, and 2 mM CaCl_2_ (pH 7.35). Negative pressure was applied through the patch pipette using an amplifier-controlled, high-speed pressure clamp system (ez-gSEAL 100b Pressure Controller; Neosystem, USA). Once the cell-attached pipette made at least a 2 ΩM seal, gentle sequential suction (−20, -40, -60 mmHg; 1 minute per pressure step) was applied to mechanically activate Piezo1 channels. In some experiments, the pipette solution was supplemented with the Piezo1 inhibitor, GsMTx4 (5 μM).

### Construction of all-points histograms

All-point histograms of single-channel currents were constructed in Clampfit. Open probability was calculated exclusively from records in which at least 10 seconds of data were recorded both before and after pressure application. The number of channels in the patch was determined by fitting a multi-Gaussian curve to all-point histograms. Consecutive channel openings were assessed by analyzing at least a 15–20-second recording.

### Immunostaining

Mouse MAs were cut open and pinned down *en face* on Sylgard blocks and fixed with 4% paraformaldehyde (PFA) at room temperature for 15 minutes. Fixed arteries were washed three times with phosphate buffered saline (PBS) for 5 minutes, then permeabilized by incubating with PBS containing 0.2% Triton-X for 1 hour at room temperature on a shaker, and blocked by incubating with permeabilizing buffer containing 5% normal donkey serum (Abcam, Waltham, MA, USA) for 1 hour at room temperature. Thereafter, MAs were incubated for 1 hour at room temperature with primary antibodies against AKAP5, Cav1, Piezo1, or TRPV4. After washing three times with PBS, block-mounted MAs were incubated with secondary antibodies (1:500; Life Technologies, Carlsbad, CA, USA; Resource Table S6) for 1 hour in the dark at room temperature. MAs were then washed three times with PBS and incubated with 0.3 µM DAPI (4′,6-diamidino-2-phenylindole) for 10 minutes at room temperature to stain nuclei. Images were acquired using an Andor Dragonfly 505 spinning-disk confocal imaging system (Andor Technology, Belfast, UK) equipped with a Borealis ILE light source; Andor Fusion software; and a Leica DMi8 microscope with an ASI PZ-2300 stage with piezo positioning, a 63X objective (NA 1.47), and an iXon888 electron-multiplying charge-coupled device (EMCCD) camera. Consecutive images were taken along the z-axis at a slice thickness of 0.2 µm from the top of endothelial cells to the bottom of SMCs (marker: smooth muscle α-actin). Images were captured using excitation lasers (405, 488, 555, and 647 nm), and emitted fluorescence was captured using band-pass filters (450/50, 525/50, 600/50, 620/60).

Arteries from knockout mice or secondary antibody alone were used as negative controls. Primary antibody-specific immunostaining was reduced in secondary antibody control and knockout mice control groups.

### *In situ* Proximity Ligation Assay (PLA)

PLA procedures were performed using a Duolink In Situ Orange Starter Kit, Mouse/Rabbit (DUO92102; Sigma-Aldrich), according to the manufacturer’s instructions. Briefly, MAs were isolated and pinned down *en face* on a Sylgard block, then fixed in 4% PFA for 15 minutes, washed three times with PBS, and incubated in a PBS/0.2% Triton X-100 solution for 20 minutes at room temperature. After incubating with the manufacturer-provided blocking buffer at 37°C for 1 hour, block-mounted arteries were washed once with PBS and incubated overnight with primary antibodies—one rabbit antibody and mouse antibody for each protein pair—at 4°C. The following day, blocks were washed three times with wash buffer A and incubated with PLA probe solution (40 µL/block) for 1 hour at 37°C in a preheated humidified chamber.m Blocks were washed three times for 5 minutes each with buffer A and incubated with ligation solution for 30 minutes at 37°C. This was followed by three more washes with buffer A and incubation with amplification solution for 100 minutes at 37°C. Amplification steps were performed in a dark room protected from direct light exposure. The blocks were then washed three times with buffer B for 5 minutes each and incubated with 0.3 µM DAPI nuclear stain (D1306; Invitrogen) for 10 minutes at room temperature in the dark. PLA images were acquired using an Andor Revolution WD (with Borealis) spinning-disk confocal imaging system (Andor Technology), comprising an upright Nikon microscope with a 60x water-dipping objective (numerical aperture, 1.0) and an electron-multiplying charge-coupled device camera (iXon888). Images were captured using excitation lasers (405 and 647 nm), and emitted fluorescence was captured using band-pass filters (450/50 and 620/60). Images were acquired along the z-axis at a slice thickness of 0.2 μm from the top surface of endothelial cells to the bottom surface where they contact SMCs. PLA signals were analyzed using Imaris 9.3 software (Oxford Instruments), which automatically detects the number of nuclei and the number of PLA puncta using surface and spots modules, respectively. The number of PLA puncta was then divided by the number of nuclei to obtain PLA puncta per cell.

### SMLM (single-molecule localization microscopy) with spectral demixing for simultaneous multi-color imaging

A dSTORM imaging system (Abbelight SAFe360; Paris, France), equipped with an automatic TIRF (total internal reflection fluorescence) module and an inverted microscope (DMi8; Leica, Wetzlar, Germany), was used to acquire super-resolution images. The microscope was configured with a 100x HC Plan-Apochromat TIRF objective (numerical aperture, 1.47; Leica), and two Hamamatsu ORCA-Fusion BT Digital CMOS cameras (to simultaneously measure intensity ratios between transmitted and reflected cameras, as described below). Images were acquired and analyzed using Neo acquisition and analysis software from Abbelight. Freshly isolated cells from mouse MAs and human skeletal muscle arteries were fixed with 4% PFA for 10 minutes, then blocked by incubating in PBS containing 5% BSA (A-421-250; Goldbio, Saint Louis, USA) and 0.2% Triton-X-100 for 1 hour at room temperature. Thereafter, cells were incubated overnight at 4°C with the primary antibodies, anti-BK, anti-PKCα and anti-TRPV4, diluted 1:100 in blocking solution. Cells were washed three times with PBS containing Tween 0.2% and incubated at room temperature for 1 hour in the dark with the following secondary antibodies (diluted 1:500 in blocking solution): CF568-conjugated donkey anti-rabbit, CF680-conjugated donkey anti-goat, and AF647-conjugated donkey anti-mouse. Cells were washed three times with PBS containing 0.1% Tween-20 and once with PBS before imaging. The primary antibodies used for cells from human subjects were anti-Piezo1 and anti-Cav1, diluted 1:100 in blocking solution. Secondary antibodies were CF680-conjugated donkey anti-rabbit and AF647-conjugated donkey anti-mouse. All detailed information of primary antibodies and secondary antibodies is listed in Table S6. Cells treated with secondary antibodies only were used for negative control experiments. For imaging, cells were embedded in super-resolution buffer solution (Smart kit; Abbelight) with microspheres (0.1 μm, T7279; Life Technologies) for alignment. All images were acquired in TIRF mode.

Automated spectral demixing involves the use of a single excitation laser to simultaneously excite two spectrally close fluorophores, with the emitted fluorescence separated into two distinct detection channels using a dichroic beam splitter. Spectral limits are computed based on automated fitting of the two populations, and ranges are set to allow an estimated crosstalk maximum of 2%. This technique eliminates chromatic aberration issues and drift associated with sequential color acquisitions (Figure S7). For image acquisition from mouse cells, we used simultaneous imaging of two proteins (BK and PKCα), followed by sequential imaging of the third protein (TRPV4). For image acquisition of cells from human subjects, we used simultaneous imaging of two proteins (Cav1 and Piezo1). The far-red fluorophores (mouse cells: AF647 and CF680 for BK and PKCα, respectively; human cells: AF647 and CF680 for Cav1 and Piezo1, respectively) were excited using a 640 nm (500 mW) laser. The emitted light was spectrally separated into two channels (transmitted and reflected) using a long-pass dichroic beam splitter (700 nm; Abbelight), collected with a 700/75 emission filter, and recorded simultaneously on two different cameras. Sequential imaging of TRPV4 was performed using a 561 nm (200 mW) excitation laser, and emitted fluorescence was collected with a 600/50 nm emission filter.(*23*) The demixing ratio for each single molecule detected is calculated as follows:

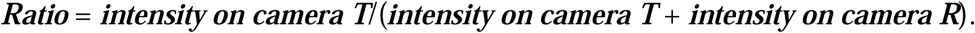

Optimal image results were achieved by combining corrections and background removal using NEO analysis software (version 40; Abbelight). Localization maps were reconstructed from 20,000 images, acquired at a rate of 20 frames per second. The first 1000 images were excluded from reconstruction to account for the time required to stabilize the photoswitching of probes. Software drift and alignment corrections were applied using model-based cross-correlation. The number of molecules/μm^2^ was assessed by subtracting localizations in the background from those at the cell membrane using the Spatial module of NEO analysis software (version 40; Abbelight).

Cluster analysis was performed using a DBSCAN (density-based spatial clustering of applications with noise) algorithm implemented in NEO Analysis software (version 40; Abbelight). This approach identifies clusters based on the spatial density of detected localizations. A minimum of 10 points within a radius of 80 nm was used to define a cluster for BK and PKCα, and a minimum of 25 points within a radius of 40 nm was used to define a cluster for TRPV4.

Colocalization analysis of dSTORM images was performed using IMARIS software (version 9.9; Andor Technology Inc., Concord, MA, USA). Spot objects were created in IMARIS using the automatic spot detection algorithm, which is based on fluorescence intensity. The intensity threshold was set manually to exclude background signal while retaining true positive puncta. These spots were then used in quantitative colocalization analysis using the Imaris Colocalization module. The percentage of BK and PKCα that colocalized with TRPV4 within 50 nm was determined from automated counts of the total number of BK, PKCα, and TRPV4 puncta in each field of view.

### Chronic Ang II infusion in mice

Mice were infused with Ang II (1 μg/kg/min) or 0.9% saline for 14 days (3). Briefly, mice were anesthetized with isoflurane (1.5%) and implanted with an osmotic micropump (Alzet Model 1004; DURECT Corporation, Cupertino, CA, USA) containing Ang II or saline. Blood pressure was monitored in conscious mice using radiotelemetry, as described above.

### Statistical analysis

Results are presented as means ± standard error of the mean (SEM). Experimental subjects and pharmacological treatments were randomized. Sample sizes were determined through power analysis using GLIMMPSE software to ensure a statistical power greater than 0.8 (a = 0.05) for detecting a change of more than 20%. Data were derived from a minimum of three mice across at least two independent experimental cohorts. N = 1 was defined as one artery in imaging and pressure myography experiments, and one subject for studies in human tissues. The normality of data distributions was assessed using Shapiro-Wilk and Kolmogorov-Smirnov tests. Non-normally distributed data were analyzed using the Mann-Whitney U test for comparisons between two groups, or the Kruskal-Wallis test followed by Dunn’s *post hoc* test for multiple comparisons. For normally distributed data, comparisons involving more than two groups were analyzed using one-way or two-way analysis of variance (ANOVA) followed by Tukey’s post hoc test for multiple comparisons. Statistical analyses were performed using GraphPad Prism 10.6 (San Diego, CA, USA). A P-value < 0.05 was considered statistically significant; “ns” denotes “not significant” (P > 0.05). Figures were prepared using CorelDraw Graphics Suite 2025 (Ottawa, ON, Canada).

## Acknowledgments

The authors thank Dr. Gary Owens for *Myh11-*Cre^ERT2^RAD mice and A. Impagliazzo for the illustrations.

## Author contributions

Conceptualization: YLC, SKS

Methodology: YLC, GCG, DCN, SBA, RM, RTK, SKS Investigation: YLC, MK, FA, YT, ZD, KK, LH, EDC, SV, SSK, SKS

Visualization: YLC, MK, FA, YT, LH, SSK, SKS Supervision: SKS

Writing—original draft: YLC, SKS

Writing—review & editing: YLC, MK, KK, RTK, SKS

## Data and materials availability

All data are available in the main text or the supplementary materials. The data, analytical methods, and materials are available from the corresponding author on reasonable request.

