## Supplemental Figures and Tables for "AKAP5 and Caveolin-1 organize opposing nanodomains that regulate smooth muscle contraction and blood pressure"

Yen-Lin Chen *et al.*

**This PDF file includes:**

Figs. S1 to S12

Tables S1 to S6

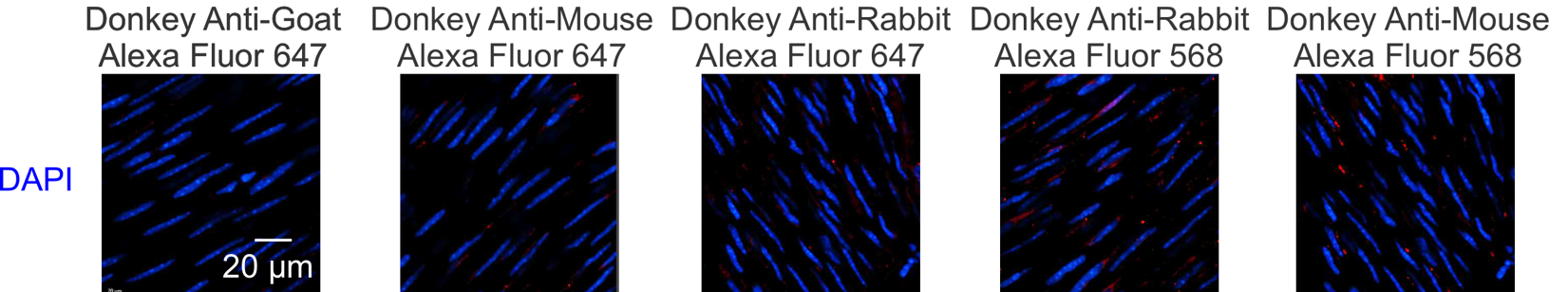

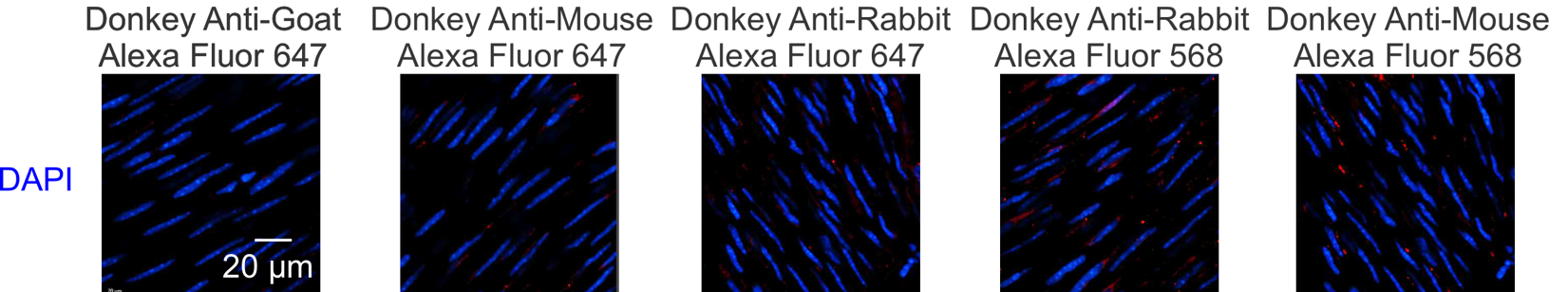

Fig. S1.

**Negative control for immunostaining.** Images of en face preparations of MAs from control mice exposed to secondary antibody only.

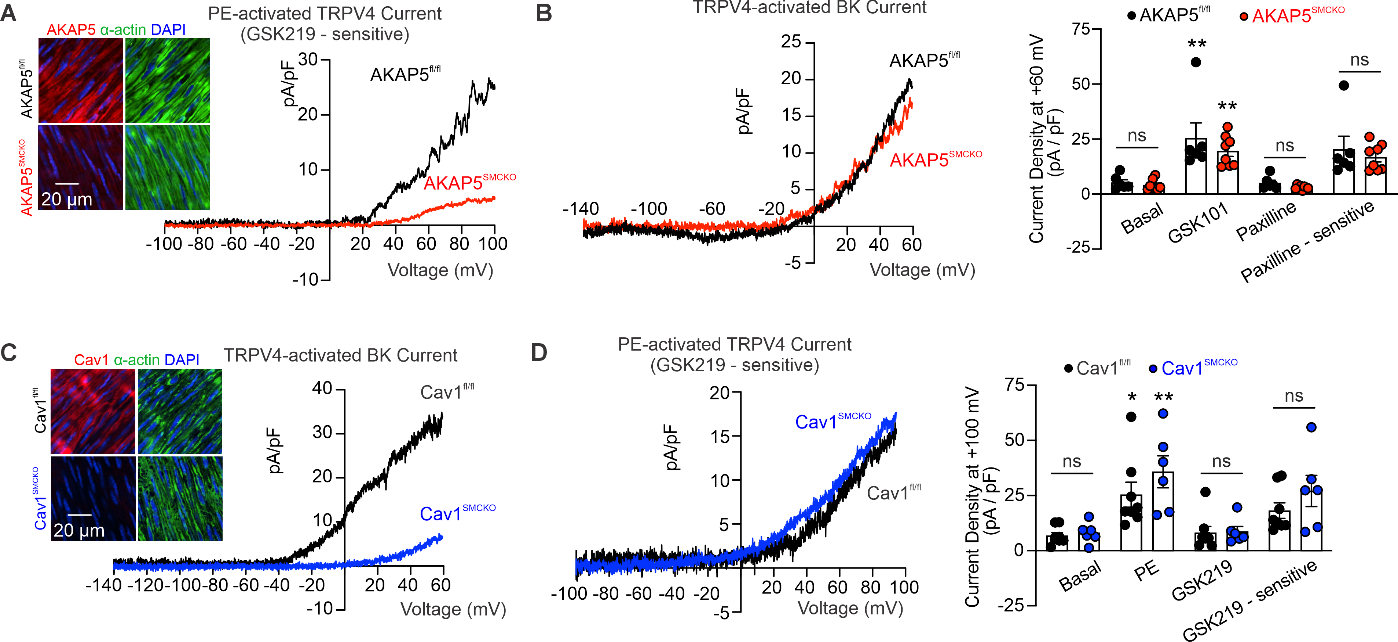

Fig. S2.

**SMCs from female mice also show AKAP5-dependent** α**_1_AR–TRPV4_SMC_ and Cav1-based TRPV4–BK channel signaling.** (A) *Inset*, representative images of AKAP5 (red), a-actin (green; SMC marker), and nuclear staining (DAPI; blue) in *en face* preparations of MAs from female AKAP5^fl/fl^ and AKAP5^SMCKO^ mice. Representative traces of ionic currents through TRPV4_SMC_ channels in SMCs from MAs of female AKAP5^fl/fl^ and AKAP5^SMCKO^ mice, presented as phenylephrine (PE)-activated GSK219-sensitive currents. (B) *Left*, representative traces of ionic currents through GSK101-activated BK channels in isolated SMCs from MAs of female AKAP5^fl/fl^ mice and AKAP5^SMCKO^ mice. *Right*, averaged outward currents at +60 mV in isolated SMCs from MAs of female AKAP5^fl/fl^ (n = 6) and AKAP5^SMCKO^ (n = 8) mice before and after administration of GSK101 (30 nM) or paxilline (1 μM) (**P < 0.01 vs. basal; ns, not significant; 2-way ANOVA). (C) *Inset*, representative images of Cav1 (red), a-actin (green; SMC marker), and nuclear staining (DAPI; blue) in *en face* preparations of MAs from female Cav1^fl/fl^ and Cav1^SMCKO^ mice. Representative traces of ionic currents through GSK101-activated BK channels in SMCs from MAs of female Cav1^fl/fl^ and Cav1^SMCKO^ mice. (D) *Left*, representative traces of ionic currents through TRPV4 channels in isolated SMCs from MAs of female Cav1^fl/fl^ and female Cav1^SMCKO^ mice, presented as PE-activated, GSK219-sensitive currents. Experiments were performed in the presence of ruthenium red (1 μM) to block Ca^2+^ entry at negative voltages. *Right*, averaged outward currents in isolated SMCs from MAs of female Cav1^fl/fl^ (n = 8) and Cav1^SMCKO^ mice (n = 6) at +100 mV before and after administration of PE (1 μM) or GSK219 (100 nM) (*P < 0.05, **P < 0.01 vs. basal; ns, not significant; 2-way ANOVA).

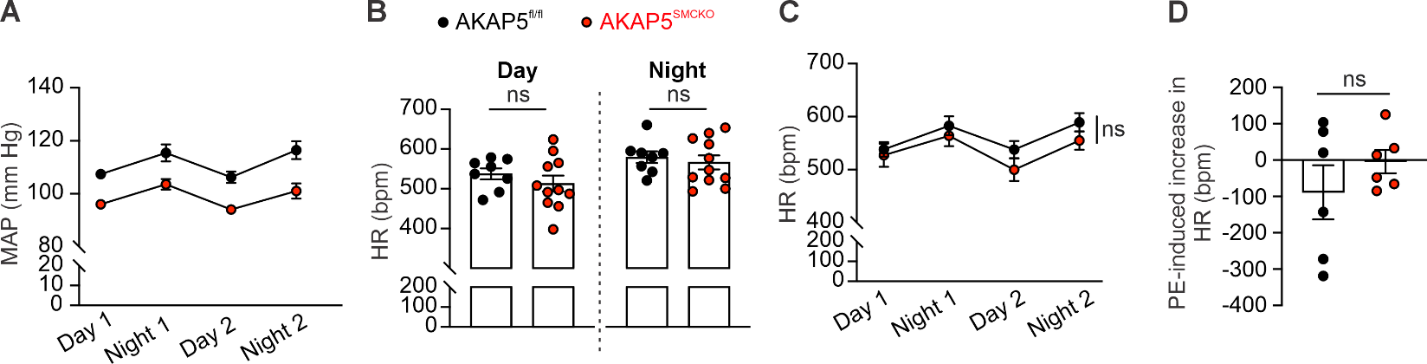

Fig. S3.

(A) Resting mean arterial pressure (MAP) in AKAP5^fl/fl^ (n = 8) and AKAP5^SMCKO^ (n = 11) mice recorded over 2 days and 2 nights. (B) Quantification of heart rate (HR) during daytime and nighttime in AKAP5^fl/fl^ (n = 8) and AKAP5^SMCKO^ (n = 11) mice (ns, not significant; unpaired t-test). (C) HR in AKAP5^fl/fl^ (n = 8) and AKAP5^SMCKO^ (n = 11) mice recorded over 2 days and 2 nights (ns, not significant; 2-way ANOVA). (D) Change in HR after a single bolus injection of phenylephrine (10 mg/kg i.p.) (n = 6; unpaired t-test).

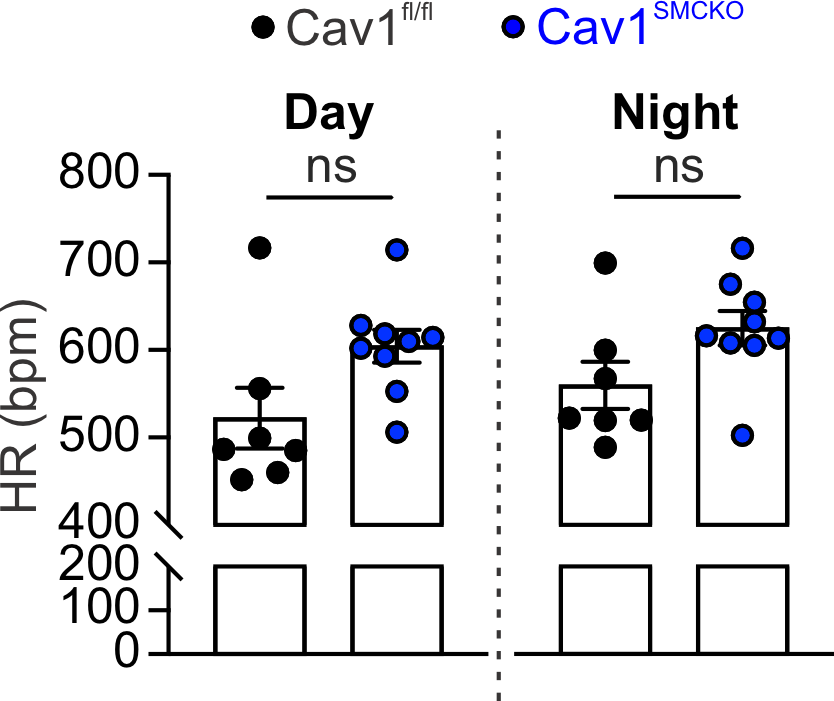

**Fig. S4.**

Quantification of heart rate (HR) during daytime and nighttime in Cav1^fl/fl^ (n = 7) and Cav1^SMCKO^ (n = 9) mice (ns, not significant; unpaired t-test).

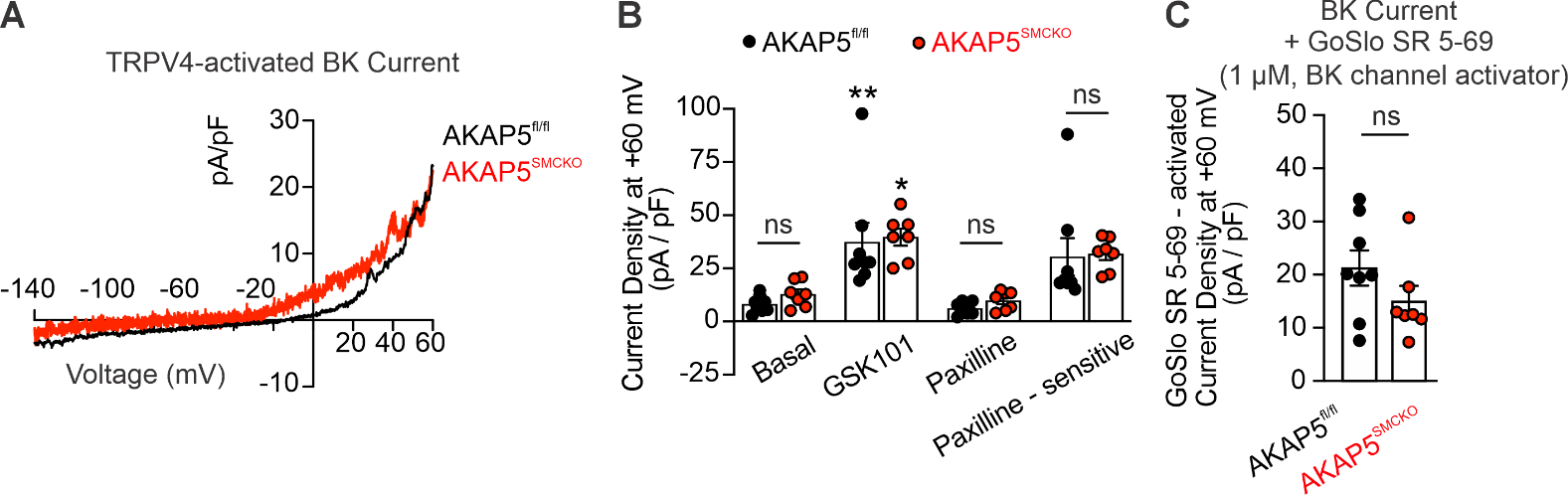

**Fig. S5.**

(A) Representative traces of ionic currents through BK channels in freshly isolated SMCs from MAs of AKAP5^fl/fl^ and AKAP5^SMCKO^ mice, recorded in the whole-cell patch-clamp configuration. (B) Outward currents at +60 mV in SMCs from MAs of AKAP5^fl/fl^ (n = 7) and AKAP5^SMCKO^ (n = 7) mice before and after treatment with GSK101 (30 nM) or GSK101+paxilline (1 μM) (*P < 0.05, **P < 0.01 vs. basal; ns, not significant; 2-way ANOVA). (C) GoSlo SR 5-69 (BK channel activator; 1 μM)-activated outward currents (GoSlo SR 5-69 minus baseline current) in SMCs isolated from MAs of AKAP5^fl/fl^ (n = 8) and AKAP5^SMCKO^ (n = 7) mice (ns, not significant; unpaired t-test).

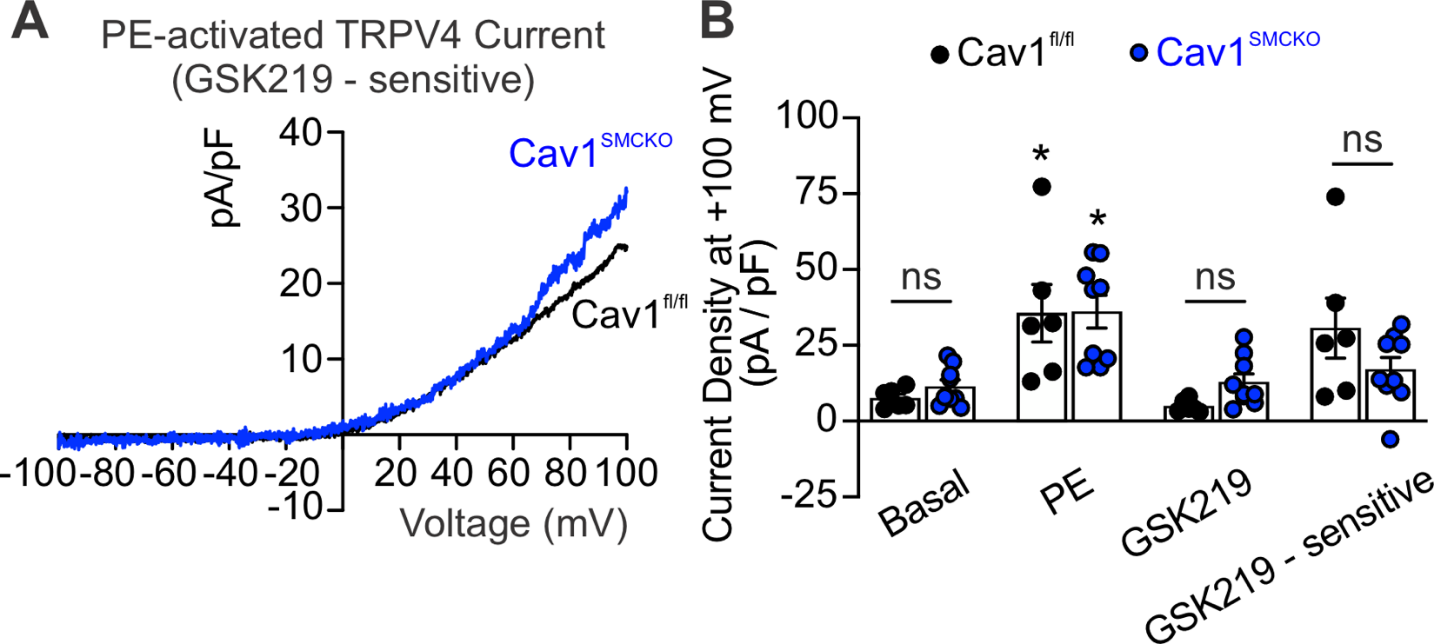

**Fig. S6.**

(A) Representative traces of ionic currents through TRPV4 channels in SMCs from MAs of Cav1^fl/fl^ mice and Cav1^SMCKO^ mice, recorded in the whole-cell patch-clamp configuration and presented as GSK219-sensitive currents. Experiments were performed in the presence of ruthenium red (1 μM) to block Ca^2+^ entry at negative voltages and subsequent activation of BK channels. (B) Outward currents in SMCs from MAs of Cav1^fl/fl^ (n = 6) and Cav1^SMCKO^(n = 9) mice before and after the addition of PE (1 μM) or PE+GSK219 (100 nM) (*P < 0.05 vs. basal; ns, not significant; 2-way ANOVA).

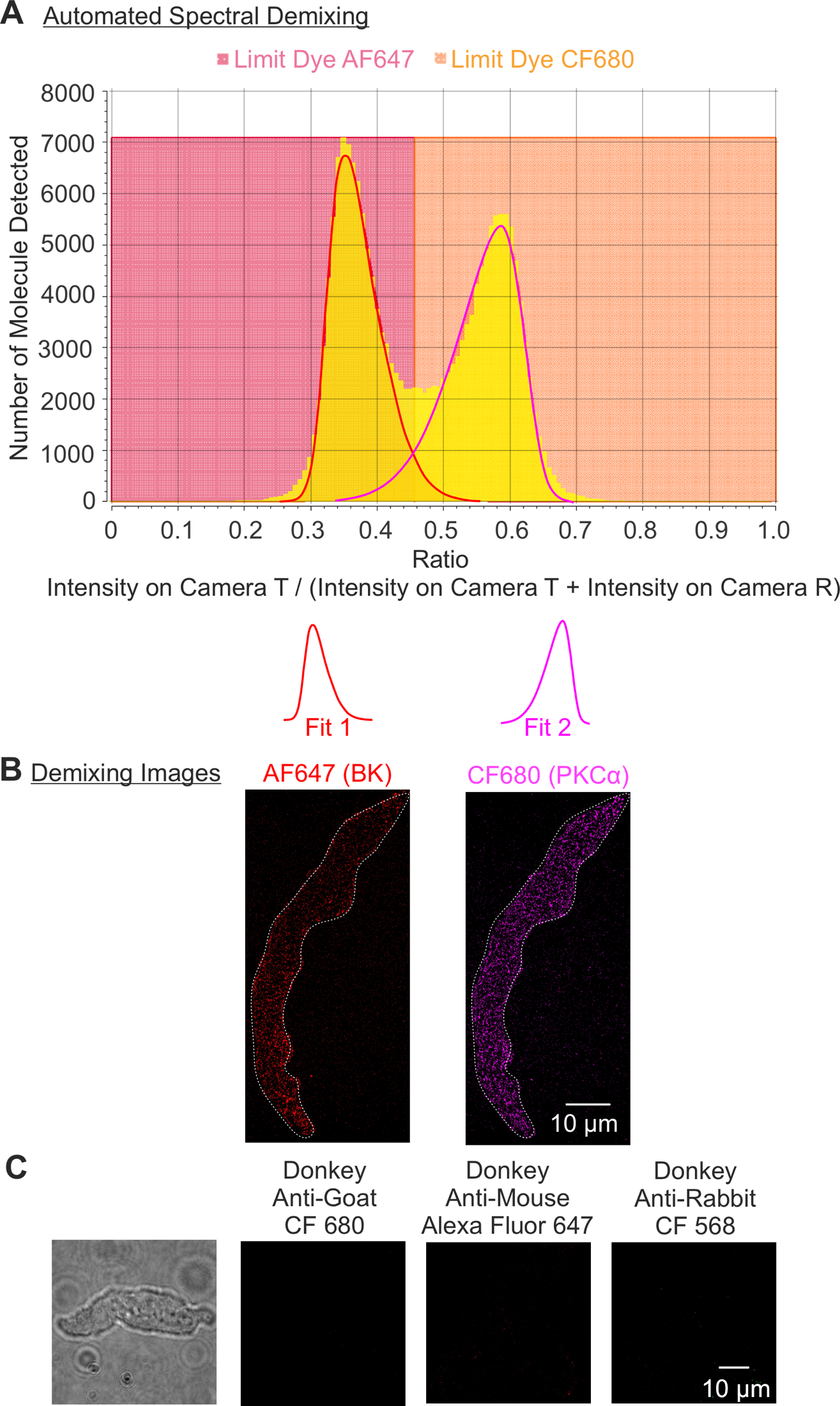

**Fig. S7.**

**SMLM technology with spectral demixing.** (A) On the graph, the vertical lines indicate the ratio limits used for spectral demixing. Ratio limits are computed based on automated fitting of the two populations (corresponding to Dye AF647 and Dye CF680), and ranges are set to allow for an estimated maximum crosstalk. (B) Representative TIRF-SMLM images acquired using spectral demixing for AF647 (BK channel) and CF680 (PKCα) localizations at the cell membrane of SMCs from mouse MAs. (C) Negative control for TIRF-SMLM images exposed to secondary antibody alone in SMCs from mouse MAs.

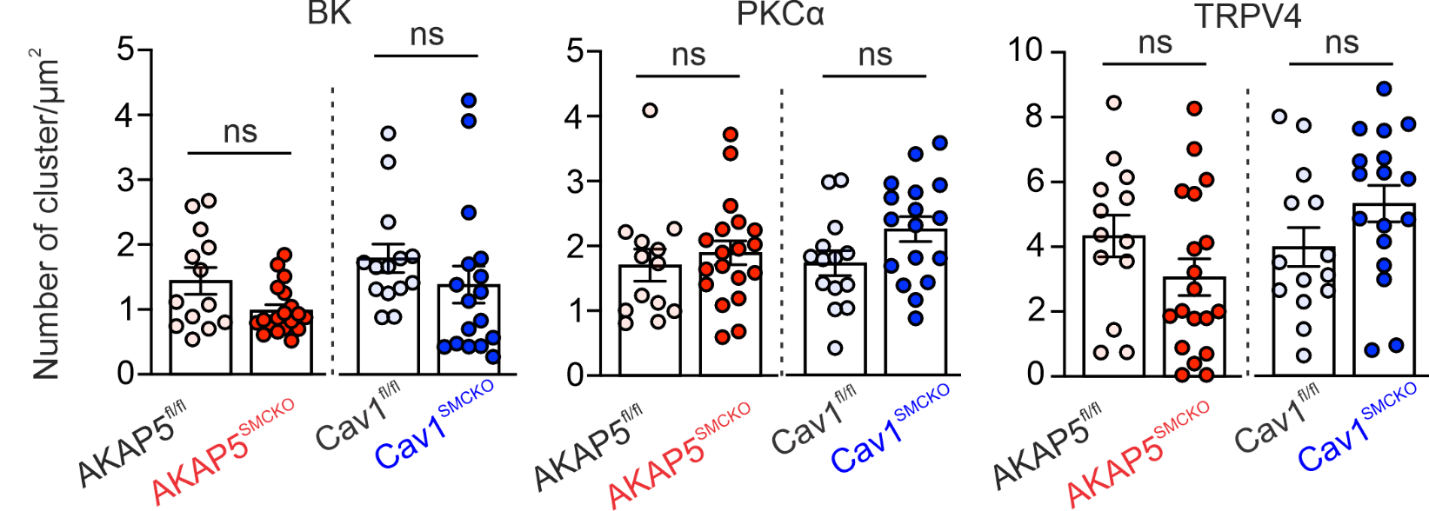

**Fig. S8.**

Cluster density of BK, PKCα, and TRPV4 channels at the cell plasma membrane of SMCs from AKAP5^fl/fl^ (n = 8), AKAP5^SMCKO^ (n = 10), Cav1^fl/fl^ (n = 8), and Cav1^SMCKO^ (n = 10) mice (1–2 cells/mouse; ns, not significant; unpaired t-test).

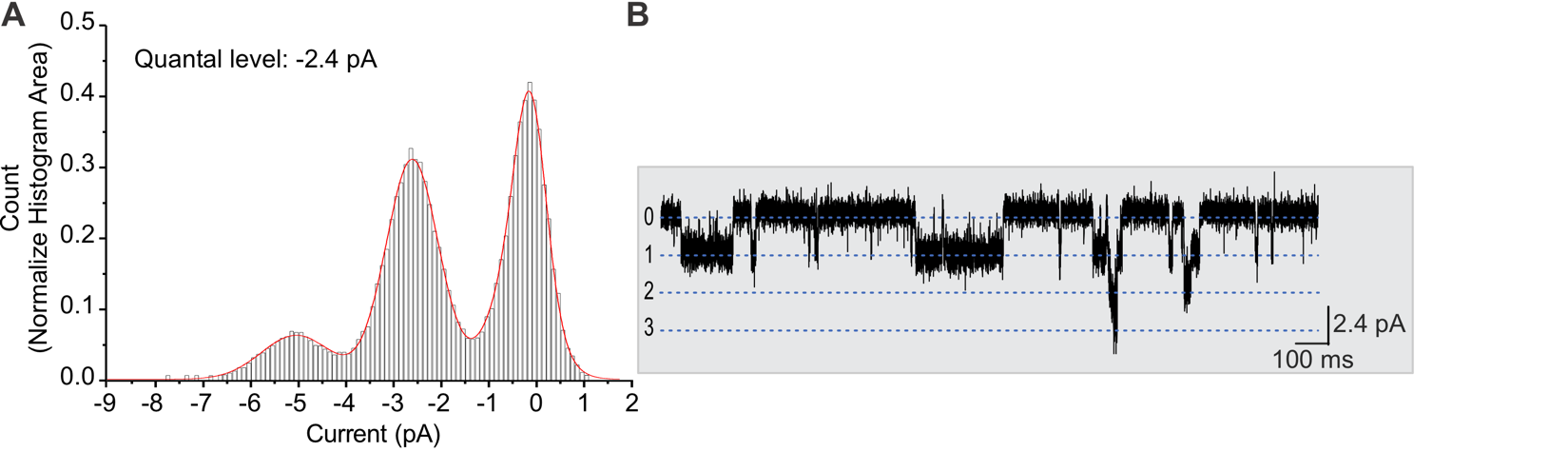

**Fig. S9.**

(A) All-points amplitude histogram of inward current traces from freshly isolated SMCs from Control (Piezo1^fl/fl^) mice. The amplitude histogram was fit to a multi-Gaussian curve using Clampfit. (B) Traces showing an example of inward Piezo1 currents in SMCs. Dotted blue lines represent quantal levels (single-channel amplitudes) determined from the all-points histogram in A.

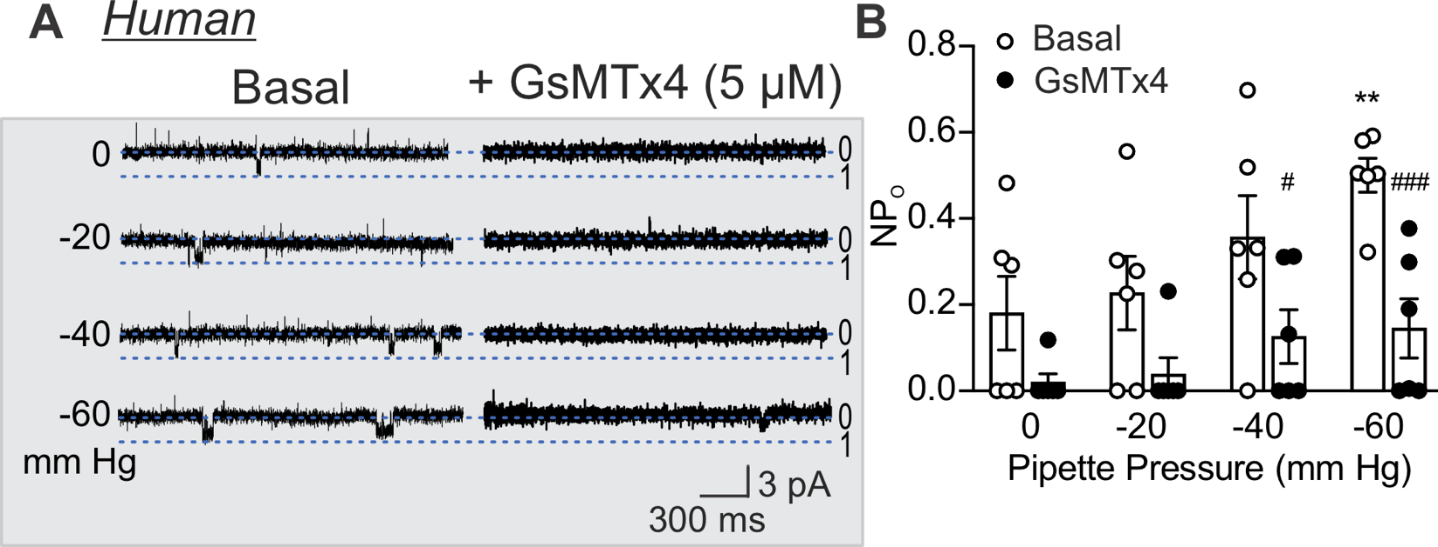

**Fig. S10.**

(A) Representative traces of inward currents in freshly isolated SMCs from a normotensive individual in a cell-attached patch. Gradually increasing suction elicited inward currents that were inhibited by the non-specific Piezo1 inhibitor, GsMTx4 (5 μM). Dotted blue lines represent quantal levels (single-channel amplitudes) determined from all-points histograms. (B) Channel activity (NP_O_, where N is the number of channels and P_O_ is the open state probability) with gradually increased suction (0, -20, -40, and -60 mm Hg) in SMCs isolated from skeletal muscle arteries of normotensive individuals (n = 6) (**P < 0.01 vs. 0 mm Hg pipette pressure; ^#^P < 0.05, ^###^P < 0.001 vs. basal; ns, not significant; 2-way ANOVA).

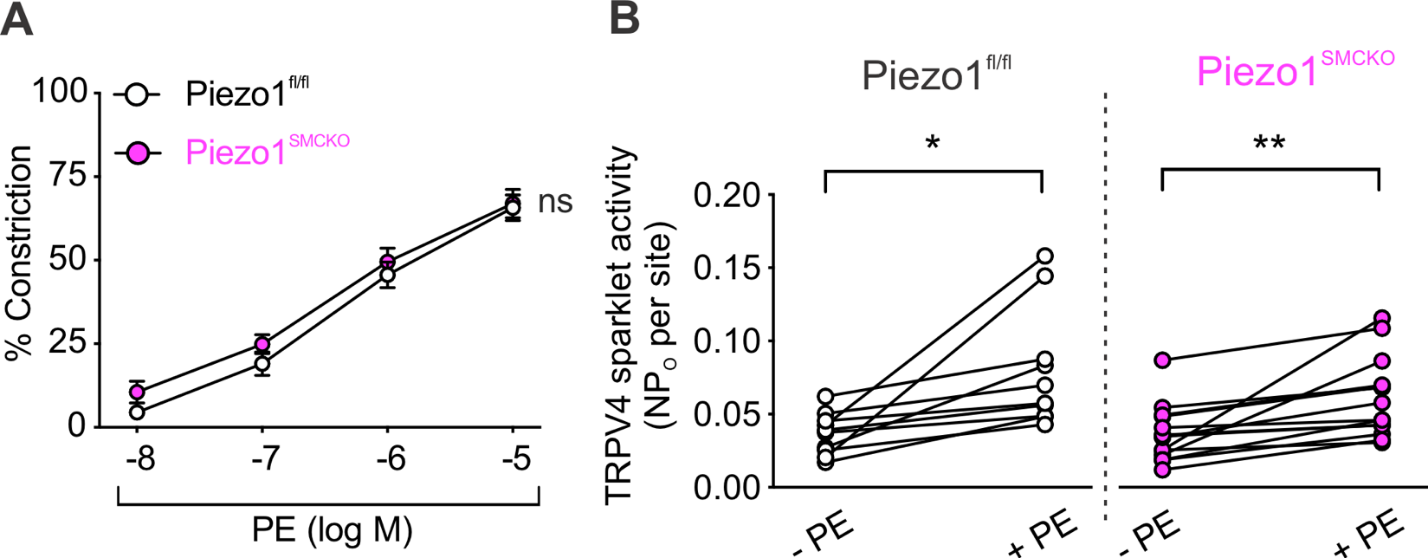

**Fig. S11.**

(A) Pressure myography results showing phenylephrine (PE)-induced constriction of small MAs from Piezo1^fl/fl^ (n = 6) and Piezo1^SMCKO^ (n = 8) mice (ns, not significant; 2-way ANOVA). (B) TRPV4_SMC_ sparklet activity (NP_O_ per site) in MAs from Piezo1^fl/fl^ (n = 10) and Piezo1^SMCKO^ (n = 14) mice. Pressurized MAs were pretreated with cyclopiazonic acid (CPA; 20 μM), nifedipine (1 μM), and GSK101 (30 nM), and then treated with PE (1 μM; *P < 0.05, **P < 0.01 vs. without PE; paired t-test).

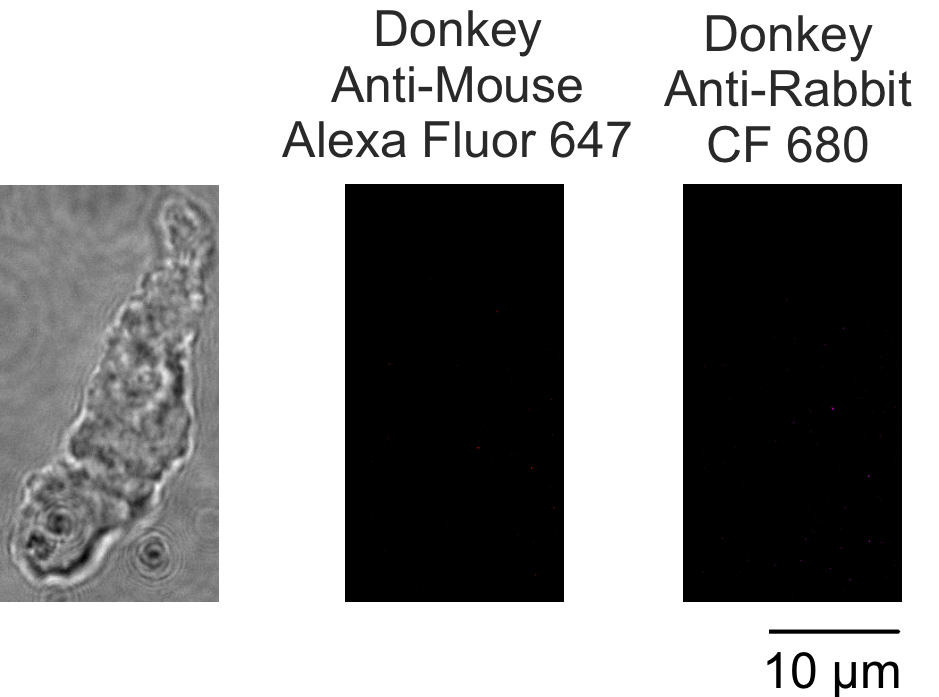

**Fig. S12.**

*Left*, snapshot of an entire SMC freshly isolated from a human skeletal muscle artery. Representative TIRF-SMLM images were acquired using spectral demixing for secondary antibody alone groups (donkey anti-mouse AF647 (*Middle*) or donkey anti-rabbit CF680 (*Right*)) in SMCs from a human subject.

**Table S1: Animals**

| **Species** | **Source** | **Background**  **Strain** | **ID/URL** |
| --- | --- | --- | --- |
| *Cav1^tm1Tt^* | Dr. Richard Minshall | C57Bl6/J | https://doi.org/10.1016/S0002-9440(10)63920-X  https://www.informatics.jax.org/allele/MGI:2654177 |
| *Akap5^tm1Jsco^* | The Jackson Laboratory | C57Bl6/J | https://www.informatics.jax.org/allele/MGI:3809936 |
| *Piezo1^tm2.1Apat^* | The Jackson Laboratory | C57Bl6/J | https://www.jax.org/strain/029213 https://www.alliancegenome.org/allele/MGI:5766345 |
| *Trpv4^tm1.1Ldtk^* | Dr. Wolfgang Liedtke | C57Bl6/J | https://doi.org/10.1073/pnas.1312933110 https://www.informatics.jax.org/allele/MGI:5544606 |
| *C57BL/6-Tg(Myh11-cre/ERT2)F31Gko/J* | Dr. Gary  Owens | C57Bl6/J | https://www.ahajournals.org/doi/10.1161/ATVBAHA.  122.318160  https://www.jax.org/strain/037658 |

**Table S2: Transnetyx genotyping probe ID**

| **Target gene** | **Probe ID** |
| --- | --- |
| *AKAP5^fl/fl^* | Akap5-3 WT, Akap3-5 FL |
| *Piezo1^fl/fl^* | Piezo1-2 WT, Piezo1-2 MD, Piezo1-2 tm1d |
| *Myh11Cre^ERT2^RAD* | Myh11-4 TG |

**Table S3: In-house genotyping primers**

| **Target gene** | **Primer sequences** |
| --- | --- |
| *Cav1^fl/fl^* | F: TTCTGTGTGCAAGCCTTTCC  R: GTGTGCGCGTCATACACTTG |
| *Trpv4^fl/fl^* | F: TTTCAGACACAAGCTTTCTAGGGTA  R: GCATCGGCTCAGCACAAAACC  R1: AAGGAGGGAGAGTGAAGATTCATTT |
| *Cav1^fl/fl^* | F: TTCTGTGTGCAAGCCTTTCC  R: GTGTGCGCGTCATACACTTG |
| *Piezo1^fl/fl^* | F: GCCTAGATTCACCTGGCTTC  R: GCTCTTAACCATTGAGCCATCT |
| *Myh11CreRad* | F: TTACCTTGGATTGCCTGGGTTTC  R: GCGAACCTCATCACTCGTTGC |

**Table S4: Chemicals/Kit/others**

| **Description** | **Catalog / Source** | **URL** |
| --- | --- | --- |
| NaCl | S7653 / Millipore Sigma | http://sigmaaldrich.com/US/en/product/sial/s7653?srsltid=AfmBOorrWtS6S17Qgd3AQNgCk2ES5vOI1Dv_gc5UezZnp1AZIQJVhX8F |
| KCl | 9541 / Millipore Sigma | https://www.sigmaaldrich.com/US/en/product/sigma/p9541 |
| KH_2_PO_4_ | P5655 / Millipore Sigma | https://www.sigmaaldrich.com/US/en/product/sigma/p5655 |
| CaCl_2_ dihydrate | C8106 / Millipore Sigma | https://www.sigmaaldrich.com/US/en/product/sial/c8106 |
| NaHCO3 | BP328500/Fisher Scientific | https://www.fishersci.com/shop/products/sodium-bicarbonate-fine-white-powder-fisher-bioreagents/BP328500 |
| MgCl_2_ hexahydrate | M2670 / Millipore Sigma | https://www.sigmaaldrich.com/US/en/product/sial/m2670 |
| Dextrose | D9434 / Millipore Sigma | https://www.sigmaaldrich.com/US/en/product/sigma/d9434 |
| HEPES | H3375 / Millipore Sigma | https://www.sigmaaldrich.com/US/en/product/sigma/h3375 |
| Papain from papaya latex | P4762 / Millipore Sigma | https://www.sigmaaldrich.com/US/en/product/sigma/p4762 |
| DL-Dithiothreitol | D0632 / Millipore Sigma | https://www.sigmaaldrich.com/US/en/product/sial/d0632 |
| Collagenase type IV | LS004186 / Worthington Biochemical Corporation | https://www.worthington-biochem.com/products/collagenase?p=122#product-122 |
| Elastase | LS006365 / Worthington Biochemical Corporation | https://www.worthington-biochem.com/products/elastase?v=564 |
| Soybean trypsin inhibitor | 10109886001 / Millipore Sigma | https://www.sigmaaldrich.com/US/en/product/roche/10109886001?srsltid=AfmBOopIuro_lW4bfZ3cCAYk4RbrnPnWEWaQKa2y7 |
| Yoda1 | 21904 / Cayman Chemical | https://www.caymanchem.com/product/21904/yoda1?srsltid=AfmBOopuBf11SZrn_uTJtL3DOagcYf91Bxa_kWX-tdJFDNk1ox7CGLwq |
| GsMTx4 | HY-P1410 / Medchemexpress | https://www.medchemexpress.com/gsmtx4.html?utm_source=google&utm_medium=CPC&utm_campaign=USHY-P1410GsMTx4&gad_source=1&gad_campaignid=18091196136&gbraid=0AAAAADnIT_YA0xxS9jp4FNt75OYUuauXG&gclid=Cj0KCQjw6bfHBhDNARIsAIGsqLiVxRQ_VGJZNihlXlBsYCgRJS6swzb-ydKLm0MGbIKC7SiiQyYP5DEaAogaEALw_wcB |
| Sodium Pentobarbital | 25021-676-20 / Diamondback Drugs | https://mms.mckesson.com/product/1273371/Hikma-Pharmaceuticals-USA-24201001020 |
| DAPI | D1306 / Invitrogen | https://www.thermofisher.com/order/catalog/product/D1306 |
| Goat serum | SER005 / Neuromics | https://www.neuromics.com/SER005 |
| Donkey serum | SER004 / Neuromics | https://www.neuromics.com/SER004 |
| Bovine serum albumin (BSA) | A2058 / Millipore Sigma | https://www.sigmaaldrich.com/US/en/product/sigma/a2058?srsltid=AfmBOoohyNM5sKI8SlKO7wYC6aRHWPfvgTI6ZNI6rYweTmWmQH92nE7M |
| Bovine serum albumin | A-421 / Gold Biotechnology | https://www.goldbio.com/products/bovine-serum-albumin-bsa-fraction-v-fatty-acid-free-for-tissue-culture |
| Normal chow diet | Teklad LM-485, 7912, Envigo | https://www.inotiv.com/rodent-traditional-natural-ingredient-diets |
| Tamoxifen diet Envigo Diet | TD130856 / Envigo diet | https://www.inotiv.com/tamoxifen-custom-diets |
| Cyclopiazonic acid (CPA) | 11326 / Cayman Chemical | https://www.caymanchem.com/product/11326/cyclopiazonic-acid?srsltid=AfmBOorSpCitLx77uCxIee8GKYMLiq5HYLe53pU8fTwPwKf2kHPT18cI |
| Ruthenium Red | Cat. No. 1439 | https://www.tocris.com/products/ruthenium-red_1439?gclsrc=aw.ds&gad_source=1&gad_campaignid=797771454&gbraid=0AAAAADRLDxSvAsCKQNld_eyh-BVn7HjSO&gclid=CjwKCAiAv5bMBhAIEiwAqP9GuKvJR3iavnM22XZud3uEJK2C7rxchbjwUN74yt2Y3akufRygJIT0UhoC4wwQAvD_BwE |
| GSK2193874 | 17715 / Cayman Chemical | https://www.caymanchem.com/product/17715/gsk2193874 |
| GSK1016790A | 17289 / Cayman Chemical | https://www.caymanchem.com/product/17289/gsk1016790a?srsltid=AfmBOooM94S3RqXPRrB-iyOFdf-LDuqMNBkPsAg2tYaddv49PJ3I2HaN |
| Phenylephrine | 2838 / Tocris Bioscience | https://www.tocris.com/products/r-phenylephrine-hydrochloride_2838 |
| Gö 6976 | 2253 / Tocris Bioscience | https://www.tocris.com/products/go-6976_2253 |
| Angiotensin II | 4006473 / BACHEM | https://shop.bachem.com/product/4006473/ |
| Nifedipine | 1075 / Tocris Bioscience | https://www.tocris.com/products/nifedipine_1075 |
| Capsaicin | 92350 / Cayman Chemical | https://www.caymanchem.com/product/92350/capsaicin |
| Atropine | 12008 / Cayman Chemical | https://www.caymanchem.com/product/12008/atropine |
| Prazosin hydrochloride | 0623 / Tocris Bioscience | https://www.tocris.com/products/prazosin-hydrochloride_0623 |
| Paxilline | 2006 / Tocris Bioscience | https://www.tocris.com/products/paxilline_2006 |
| GoSlo SR 5-69 | 5814 / Tocris Bioscience | https://www.tocris.com/products/goslo-sr-5-69_5814 |
| Fluo‐4 AM | F14201 / ThermoFisher | https://www.thermofisher.com/order/catalog/product/F14201?ef_id=Cj0KCQjw6bfHBhDNARIsAIGsqLj9xY0GE_mtuSeYgedq0KOEDnptNSdbGk_4rReN_aF80FJA7p44xMQaAlCGEALw_wcB:G:s&s_kwcid=AL!3652!3!447292198772!!!g!!!10506731179!109642167771&cid=bid_pca_iva_r01_co_cp1359_pjt0000_bid00000_0se_gaw_dy_pur_con&gad_source=1&gad_campaignid=10506731179&gbraid=0AAAAADxi_GQ7gMmsrfTYbgQS5Q7G0W88M&gclid=Cj0KCQjw6bfHBhDNARIsAIGsqLj9xY0GE_mtuSeYgedq0KOEDnptNSdbGk_4rReN_aF80FJA7p44xMQaAlCGEALw_wcB |
| AAT Bioquest Pluronic F-127 | AATB-20053 / Millipore Sigma | https://www.sigmaaldrich.com/US/en/product/aatbioquest/aatb20053?srsltid=AfmBOoqLg8C7BC_e6vPsLxPoWXT3yDOreE5G6-At95Y3d4gB212m1SiA |
| DMSO | D4540 / Millipore Sigma | https://www.sigmaaldrich.com/US/en/product/sigma/d4540?srsltid=AfmBOooNLYCA-f9D-3r6-t5G2f-ZK-ZCUpfFoIqKyt78mRzst9V-cjG8 |
| Paraformaldehyde | 15713 / Electron Microscopy Sciences | https://www.emsdiasum.com/paraformaldehyde-20-aqueous-sol-em-grade |
| PLA-DUO92102 | DUO92102-1KT / Millipore Sigma | https://www.sigmaaldrich.com/US/en/product/sigma/duo92102?srsltid=AfmBOooyXS8dHoEZAFOUnQM38lNyKNShfMVy-NREvUgVYksoOagcYWim |
| Triton X-100 | X100 / Millipore Sigma | https://www.sigmaaldrich.com/US/en/product/sial/x100?srsltid=AfmBOooNtL6at0yqO9UoYntejj343v8Re_-HNdAQ7PN5xbwveiz53lom |
| PBS | 14190144 / ThermoFisher | https://www.thermofisher.com/order/catalog/product/14190144 |
| 100 bp DNA Ladder | N0467L / New England Biolabs | https://www.neb.com/en-us/products/n0467-quick-load-100-bp-dna-ladder?srsltid=AfmBOopS0Nklmr5S67BJVTvx6t-XMLA91DiXXZb7FlmiujFXu7DouB3P |
| Platinum SuperFi II PCR Master Mixes | 12369010 / Invitrogen | https://www.thermofisher.com/order/catalog/product/12369010?gclid=CjwKCAiAp4O8BhAkEiwAqv2UqKqz3gK6MeWIUTFEs35pbmm6CxBdAwe59E_BB_XpibO4SnIyC0y2UxoCbYMQAvD_BwE&source=google_shopping&ISO_CODE=us&LANG_CODE=en&ef_id=CjwKCAiAp4O8BhAkEiwAqv2UqKqz3gK6MeWIUTFEs35pbmm6CxBdAwe59E_BB_XpibO4SnIyC0y2UxoCbYMQAvD_BwE:G:s&s_kwcid=AL!3652!3!724104819102!!!g!2372177373295!!21983340143!166789107330&ev_chn=shop&cid=0se_gaw_25072023_BWZIOP&source=google_shopping&ISO_CODE=us&LANG_CODE=en&gad_source=1 |
| 35 mm dish-14 mm glass diameter | P35GC-1.5-14-C | https://www.mattek.com/store/p35gc-1-5-14-c-case/ |
| 35 mm dish-10 mm glass diameter | P35GC-1.5-10-C | https://www.mattek.com/store/p35gcol-1-5-10-c-case/ |

**Table S5: Clinical parameters of human subjects.**

|  | **Sex** | **Anti-HTN medicine** | **Race/ Ethnicity** | **Anti-inflamma-tory**  **medicine** | **Preop BP**  **(mm Hg)** | **Muscle**  **tissue** | **Reason for surgery** | **Co-morbidities** |
| --- | --- | --- | --- | --- | --- | --- | --- | --- |
| **Non-HTN** | F | N | White or Caucasian | N | 172/74 | Multifidus | Lumbar decompression | Breast cancer, hiatal hernia, scoliosis, spinal stenosis |
|  | F | N | African American | Y | 126/62 | Temporalis | Evacuation of a large right sided chronic subdural hematoma | congestive heart failure, heart transplant, breast cancer, type 2 diabetes, hyperlipidemia, |
|  | M | N | African American | N | 144/90 | Multifidus | Thoracic decompression | Skin cancer, Lyme disease, major depressive disorder |
|  | F | N | White or Caucasian | N | 117/63 | Multifidus | Intramedullary spinal cyst fenestration | Anemia, hypercholesterolemia, GERD, major depressive disorder |
|  | F | N | White or Caucasian | Y | 137/74 | Multifidus | Spinal deformity correction | Arthritis, scoliosis, lumbar stenosis |
|  | M | N | White or Caucasian | N | 147/91 | Temporalis | Brain tumor | GBM, Afib, hypercholesterolemia, pre-diabetes |
|  | F | N | White or Caucasian | Y | 133/60 | Temporalis | MoyaMoya | Moya Moya, stroke, carotid stenosis |
|  | F | N | White or Caucasian | N | 112/45 | Platysma | Carotid stenosis | cervical herniated disc |
|  | M | N | White or Caucasian | Y | 128/83 | Multifidus | Lumbar decompression | low back pain d/t lumbar stenosis |
|  | F | N | White or Caucasian | N | 112/75 | Multifidus | Lumbar decompression | low back pain d/t lumbar stenosis |
|  | M | N | White or Caucasian | N | 115/68 | Multifidus | Spinal deformity correction | Hypercholesterolemia, atrial fibrillation, sarcoidosis, melanoma, restless leg syndrome, sleep apnea |
|  | M | N | White | N | 98/60 | SCM | Left-sided Vagal nerve stimulator implantation for stroke motor weakness | stroke 2/2 AVM, Seizures, Cavernoma, PNET, AVM |
|  | F | N | White | Y | 133/73 | SCM | Left-sided Vagal nerve stimulator implantation for stroke motor weakness | Afib, stroke |
|  | F | N | White | Y | 111/62 | SCM | Left sided Vagal nerve stimulator implantation for stroke motor weakness | Stroke, HLD |
| **HTN** | M | Y | White or Caucasian | N | 96/59 | Multifidus | Lumbar discitis and osteomyelitis, spinal deformity correction | Anemia, C difficile colitis, CHF, CKD, GERD, Guillain-Barré, essential tremor |
|  | F | N | White or Caucasian | Y | 175/83 | Multifidus | Lumbar decompression | DM, GERD |
|  | M | Y | White or Caucasian | Y | 138/90 | Multifidus | Resection of sacral mass | Alcohol use disorder, hypercholesterolemia, psoriasis, previous stroke |
|  | M | N | White or Caucasian | N | 140/66 | Multifidus | Lumbar decompression | OSA (uses CPAP), hypercholesterolemia, Charcot-Marie Tooth Disease, atrial fibrillation |
|  | F | Y | White or Caucasian | N | 121/73 | Splenius | Cervical decompression | Hyperlipidemia, chronic fatigue, major depressive disorder, generalized anxiety disorder, ADHD, GERD, insomnia, tinnitus, and vitamin D deficiency |
|  | F | Y | White or Caucasian | Y | 141/81 | Temporalis | Meningioma resection | Hyperlipidemia, hypercalcemia, temporal lobe epilepsy, vitamin D deficiency, major depressive disorder, generalized anxiety disorder, fatty liver |
|  | F | Y | White or Caucasian | Y | 116/58 | Multifidus | Lumbar decompression | GERD, hypercholesterolemia, psoriasis, Sjogren's syndrome |
|  | M | Y | Hispanic | Y | 146/68 | Temporalis | Moya Moya | HTN, hypercholesterolemia, Moya Moya |
|  | M | N | White or Caucasian | N | 148/83 | Multifidus | Lumbar decompression | HTN, hypercholesterolemia, CKD, cirrhosis, GERD, pre-diabetes, aortic valve stenosis |
|  | F | Y | White or Caucasian | Y | 139/65 | Multifidus | Lumbar decompression | HTN, hypercholesterolemia, type II diabetes, osteoporosis, hypothyroidism, GERD |
|  | M | Y | White or Caucasian | Y | 152/72 | Multifidus | Lumbar decompression | CAD, OSA, HLD, MI, diabetes, lumbar stenosis |
|  | F | Y | White or Caucasian | Y | 150/64 | Multifidus | Lumbar decompression | HTN, hypercholesterolemia, coronary artery disease, abdominal aortic aneurysm, type II diabetes, fatty liver, OSA, osteoarthritis |
|  | M | Y | White or Caucasian | Y | 124/80 | Multifidus | Lumbar decompression | HTN, prostate cancer, polycythemia, lupus, polymyalgia rheumatica, prior spinal cord injury |
|  | M | Yes | White or Caucasian | N | 166/78 | Multifidus | Spinal deformity correction | HTN, sleep apnea, history of HIV (Dx 1999, CD4 >1000s since 2002), asthma, depression |
|  | M | Y | White or Caucasian | Y | 132/61 | Multifidus | Lumbar decompression | HTN, chronic LBP, T2DM, obesity |
|  | M | Y | White or Caucasian | Y | 119/57 | Multifidus | Lumbar decompression | HTN, CAD, DM, CHF |
|  | M | Y | White or Caucasian | Y | 108/59 | Multifidus | Lumbar fusion | COPD, CKD, Guillain-Barré syndrome, hypertension, renal cell carcinoma, OSA, diabetes |
|  | F | Y | African American | Y | 153/94 | Splenius capitus | ligation of complex dural arteriovenous fistula | Diabetes, Asthma |
|  | M | N | White | N | 145/92 | Splenius capitus | Posterior fossa decompression for cerebellar infarct | Factor V Leiden, Afib, Aortic aneurysm |
|  | M | Y | White | N | 128/81 | SCM | Left sided Vagal nerve stimulator implantation for stroke motor weakness | DVT, Afib migraine, stroke |
|  | M | Y | White | Y | 156/75 | SCM | Left sided Vagal nerve stimulator implantation for stroke motor weakness | Afib, stroke |
|  | F | Y | African American | Y | 138/80 | SCM | Left sided Vagal nerve stimulator implantation for stroke motor weakness | Stroke, diabetes, hyperlipidemia |
|  | M | Y | White | N | 156/72 | SCM | Left sided Vagal nerve stimulator implantation for stroke motor weakness | Pulmonary embolus, Diabetes type 2, stroke, renal mass |

**Table S6: Antibodies used.**

| **Target Antigen** | **Source** | **Catalog** | **Concen**  **-tration** |  |
| --- | --- | --- | --- | --- |
| Cav1 Rabbit | Abcam | ab2910 | 1:100 | https://www.abcam.com/en-us/products/primary-antibodies/caveolin-1-antibody-caveolae-marker-ab2910?srsltid=AfmBOoqNKq9xQDZwOelwLl5cvm2yTZE8QljnCRnLdTog8iuOx__zAd-7 |
| Cav1  Mouse | Novus Biologicals | NB100-615 | 1:100 | https://www.novusbio.com/products/caveolin-1-antibody-7c8_nb100-615?srsltid=AfmBOopOswubylHFHF3AJYFpUIDJaga_UERAakV4Hf0w-jeF4r-WFxfH |
| AKAP150  Mouse | Santa Cruz | sc377055 | 1:100 | https://www.scbt.com/p/akap-150-antibody-e-1?srsltid=AfmBOooMSMN_TY1vPDd0Ia96zGs2kIkEA_Ol1O6HidzVjVcua8gSy9v2 |
| TRPV4  Rabbit | Lifespan Biosciences | LS-C94498 | 1:100 | https://www.lsbio.com/antibodies/trpv4-antibody-aa100-150-if-immunofluorescence-ihc-wb-western-ls-c94498/95719 |
| TRPV4  Mouse | Millipore  Sigma | MABS466 | 1:200 | https://www.sigmaaldrich.com/US/en/product/mm/mabs466?srsltid=AfmBOoomrYP0RjGaSe1b6Y_2c0gYZODgHi633bSTsjewytZSlAgLrI_d |
| PKCα  Mouse | Santa Cruz | sc17769 | 1:100 | https://www.scbt.com/p/pkc-antibody-a-3?srsltid=AfmBOopRagOcDZg6UeYRE9UqbXb7lw6bIl5tV4D7ktYCUaObr4bpyWxc |
| PKCα  Goat | R&D Systems | AF5340 | 1:100 | https://www.rndsystems.com/products/human-mouse-rat-pkcalpha-antibody_af5340 |
| MaxiKα Antibody (B-1)  Mouse | Santa Cruz | sc374142 | 1:100 | https://www.scbt.com/p/maxikalpha-antibody-b-1?srsltid=AfmBOoooZRg8WCZgxTVYSISvhxxeiEPDXOJf-6PBHBuDBPflgi74JG_W |
| Slo1 Maxi-Potassium  Channel Antibody (L6/60) Mouse | NeuroMab | 75-022 | 1:100 | https://www.antibodiesinc.com/products/anti-slo1-maxi-k-channel-antibody-l6-60-75-022?srsltid=AfmBOoo_1k0Av5j_o1rjNYRRMjSh9z1gXjDB3YMPZOU94WI6mhfIOj8V |
| Smooth muscle α-actin (α-SMA) | Millipore  Sigma | F3777 | 1:500 | https://www.sigmaaldrich.com/US/en/product/sigma/f3777?srsltid=AfmBOoqRkAKrT43__4aGS6WalwCd7X6CCKOi9MDAj2yMTZXjf6URmj6p |
| Piezo1 Rabbit | Alomone Labs | APC-087 | 1:100 | https://www.alomone.com/p/anti-piezo1-extracellular-apc-antibody/APC-087-APC?srsltid=AfmBOoq7jVKTQ_5hadqpSXWInnx9zAGYpmehoFFDL0_blDB8d02_QUDv |
| Piezo1  Mouse | Novus Biologicals | NBP2-75617 | 1:100 | https://www.novusbio.com/products/piezo1-antibody-2-10_nbp2-75617#reviews-publications |
| Donkey Anti-Goat  Alexa Fluor 647 | Invitrogen | A21447 | 1:500 | https://www.thermofisher.com/antibody/product/Donkey-anti-Goat-IgG-H-L-Cross-Adsorbed-Secondary-Antibody-Polyclonal/A-21447 |
| Donkey  Anti-Mouse  AF647 | Invitrogen | A32787 | 1:500 | https://www.thermofisher.com/antibody/product/Donkey-anti-Mouse-IgG-H-L-Highly-Cross-Adsorbed-Secondary-Antibody-Polyclonal/A32787 |
| Donkey Anti-Rabbit  Alexa Fluor 647 | Invitrogen | A31573 | 1:500 | https://www.thermofisher.com/antibody/product/Donkey-anti-Rabbit-IgG-H-L-Highly-Cross-Adsorbed-Secondary-Antibody-Polyclonal/A-31573 |
| Donkey Anti-Rabbit  Alexa Fluor 568 | Invitrogen | A10042 | 1:500 | https://www.thermofisher.com/antibody/product/Donkey-anti-Rabbit-IgG-H-L-Highly-Cross-Adsorbed-Secondary-Antibody-Polyclonal/A10042 |
| Donkey Anti-mouse  Alexa Fluor 568 | Invitrogen | A10037 | 1:500 | https://www.thermofisher.com/antibody/product/Donkey-anti-Mouse-IgG-H-L-Highly-Cross-Adsorbed-Secondary-Antibody-Polyclonal/A10037 |
| Donkey Anti-Goat CF680 | Biotium | 20060 | 1:500 | https://biotium.com/product/donkey-anti-goat-igg-hl-highly-cross-adsorbed/?attribute_pa_conjugation=cf680 |
| Donkey Anti-Rabbit CF680 | Biotium | 20418 | 1:500 | https://biotium.com/product/donkey-anti-rabbit-igg-hl-highly-cross-adsorbed/?attribute_pa_conjugation=cf680 |
| Donkey Anti-Rabbit CF568 | Biotium | 20098 | 1:500 | <https://biotium.com/product/donkey-anti-rabbit-igg-hl-highly-cross-adsorbed/?attribute_pa_conjugation=cf568> |
